# Functional interactions between anteroventral periventricular kisspeptin neurons and gonadotropin-releasing hormone neurons in female mice

**DOI:** 10.64898/2026.02.12.705606

**Authors:** J. Rudolph Starrett, Devon Krasner, Chrystian D. Phillips, Suzanne M. Moenter

## Abstract

Kisspeptin neurons in the rostral hypothalamus are hypothesized to initiate preovulatory gonadotropin-releasing hormone (GnRH) surges by causing estradiol-dependent activation of GnRH neuron action potential firing and subsequent GnRH release. To determine if estradiol or ovarian cycle stage modulates functional connectivity in this circuit, we used optogenetics to photostimulate anteroventral-periventricular (AVPV) area kisspeptin neurons while recording electrical activity and/or evoked synaptic currents from preoptic area GnRH neurons in acutely-prepared mouse brain slices. Slices were prepared from mice in multiple hormonal states, including 2-days post ovariectomy (OVX) and OVX plus estradiol during the morning or afternoon, diestrus, proestrus and 1-week post OVX, and 6-weeks post OVX with or without 1 week of estradiol replacement. Photostimulation induced a sustained, frequency-dependent increase in GnRH neuron firing rate. This neuromodulatory-typical response was not different in diestrous vs proestrous mice but was blunted in 1-week OVX mice, suggesting ovarian steroids amplify this response. Neuromodulatory responses were infrequent in 6-week OVX mice even with 1-week of estradiol treatment. A minority of GnRH neurons exhibited a substantial and near-immediate increase in firing rate typical of fast synaptic transmission. Monosynaptic connectivity was low and stable across the hormone states tested and mediated by GABA. Interestingly, evidence of a monosynaptic connection was not a requirement for GnRH neurons to exhibit a sustained increase in firing rate, suggesting non-synaptic or volume transmission occurs in this system. Synaptic connectivity did, however, amplify the increase in firing rate observed in GnRH neurons from proestrous mice, indicating proestrous hormonal conditions can amplify this response.

**Significance statement:** Ovulation is initiated by central positive feedback effects of estradiol stimulating a surge of gonadotropin-releasing hormone (GnRH) release. Estradiol feedback is conveyed to GnRH neurons by afferents expressing estrogen receptor alpha, including kisspeptin-expressing neurons in the anteroventral periventricular (AVPV) area. To determine if endocrine milieu modulates functional interactions between AVPV kisspeptin and GnRH neurons, optogenetics was used to stimulate AVPV kisspeptin neurons while recording GnRH neuron spiking activity or synaptic currents in brain slices from ovariectomized, estradiol-treated, and ovary-intact mice. Stimulation (20Hz) increased GnRH neuron firing rate in all hormone conditions. This effect was stronger during proestrus and was further increased in GnRH neurons receiving fast-synaptic transmission. A synaptic connection was not required, however, suggesting volume transmission occurs.

## Introduction

Patterned secretion of gonadotropin-releasing hormone (GnRH) by central neurons controls fertility and is regulated by ovarian steroid feedback to the hypothalamus and pituitary. During most of the cycle, estradiol exerts negative feedback on episodic GnRH release(Goodman and Karsch, 1980; Karsch et al., 1987; Terasawa, 1994; Evans et al., 1995; Uenoyama et al., 2015; Goodman et al., 2022). In response to the high circulating estradiol levels of the late follicular phase (proestrus in rodents), however, there is a shift in estradiol feedback action from negative to positive. This changes the GnRH release pattern from strictly episodic to a prolonged surge of release lasting hours(Sarkar et al., 1976; Moenter et al., 1990, 1991; Pau et al., 1993; Levine et al., 1985). The GnRH surge causes a surge in luteinizing hormone (LH) release, which induces ovulation. While the major systemic endocrine feedback loops regulating GnRH/LH release patterns have been identified, there is comparatively less knowledge of the neural mechanisms involved.

GnRH surges are hypothesized to involve estradiol-sensitive afferents because estrogen receptor α (ERα) is required for estradiol feedback but is typically not detected in GnRH neurons(Lehman et al., 1993; Hrabovszky et al., 2000; Couse et al., 2003). Kisspeptin-expressing neurons in the hypothalamus express ERα, project to GnRH neurons and are major activators of GnRH neurons(Kinoshita et al., 2005; Wintermantel et al., 2006; Lehman et al., 2010; Cravo et al., 2011; Rønnekleiv and Kelly, 2013; Kumar et al., 2015). Loss of kisspeptin or kisspeptin signaling causes infertility(de Roux et al., 2003; Seminara et al., 2003). In mice, knockdown of ERα in rostral hypothalamic kisspeptin-expressing neurons in the anteroventral periventricular (AVPV) area blunts LH surges, indicating AVPV kisspeptin neurons serve as key intermediaries for GnRH surge induction(Wang et al., 2019; Clarkson et al., 2023). Estradiol increases kisspeptin expression in the AVPV and the action potential firing rate of AVPV kisspeptin neurons in brain slices, two mechanisms that could increase kisspeptin signaling to GnRH neurons(Smith et al., 2005; Wang et al., 2016). Consistent with this, the expression of cFos, a surrogate marker of neuronal activation, is higher in AVPV kisspeptin neurons during positive feedback, whether induced by estradiol treatment or during the preovulatory estradiol rise of the cycle(Smith et al., 2006; Wintermantel et al., 2006 p.200; Clarkson et al., 2008; Robertson et al., 2009). AVPV kisspeptin neurons also express GABAergic markers(Cravo et al., 2011). Activation of GABA_A_ receptors can excite GnRH neurons and GnRH neurons receive increased GABAergic transmission during both estradiol-induced and proestrous LH surges(DeFazio et al., 2002; Herbison and Moenter, 2011; Adams et al., 2018). This suggests AVPV kisspeptin neurons could be a source of increased GABAergic transmission to GnRH neurons during estradiol positive feedback. Estradiol thus has several actions that could poise kisspeptin neurons to increase fast-synaptic and neuromodulatory transmission to GnRH neurons, but it is not known if there is increased functional connectivity in the circuit during positive feedback.

To test if functional connectivity changes across hormonal states, we used optogenetics in acutely-prepared brain slices to photostimulate AVPV kisspeptin neurons while recording responses from GnRH neurons in the preoptic area (POA). Effects of hormonal state were examined by performing experiments in brain slices from diestrous, proestrous, and ovariectomized (OVX) mice with and without estradiol (E) replacement. In rodents, the time of estradiol positive feedback is regulated by the light/dark cycle. We thus also tested the effect of time of day in an estradiol replacement model that generates diurnal shifts in estradiol feedback action.

## Materials and Methods

Chemicals were purchased from Sigma-Aldrich unless noted

### Animals

The University of Michigan Institutional Animal Care and Use Committee approved all procedures. Adult female (80-185d) GnRH-GFP, Kiss-Cre; Kiss1-hrGFP, and Kiss-Cre;GnRH-GFP mice were used for these studies(Suter et al., 2000; Cravo et al., 2011, 2013). Mice were provided with Teklad 2916 chow (Inotiv) and water *ad libitum* and were held on a 14:10h light:dark cycle with lights on at 0300 Eastern Standard Time. Estrous cycle stage was monitored in gonad-intact mice by daily vaginal cytology from the day of stereotaxic surgery until brain slice preparation (up to 8wks). Uterine mass was measured to confirm cycle stage. Uterine mass >100□mg indicated *in vivo* exposure to a high concentration of estradiol, typical of proestrus, whereas mass <75□mg indicated exposure to low estradiol, typical of diestrus(Shim et al., 2000). One mouse with a uterine mass falling outside these ranges was excluded.

### Stereotaxic surgery

Mice were anesthetized with 1.5–2% isoflurane (Abbott). Carprofen (Zoetis, Inc., 5mg/kg, sc) was given before and 24h after surgery as an analgesic. After exposing the skull and performing a craniotomy, AVPV injections were targeted 0.2mm anterior to Bregma, 5.4mm ventral to Bregma, and ±0.2mm lateral to the center of the superior sagittal sinus. Bilateral injections of 100nL adeno-associated virus AAV) vector were made at the target coordinates at ∼5nl/min using a PLI-100A Pico-Injector (Warner Instruments). The specific vector was AAV5-Syn-FLEX-rc[ChrimsonR-tdTomato] (Addgene # 62723-AAV5) (Klapoetke et al., 2014). The injection pipette was left in place for 5min after injection to allow virus to diffuse into the brain, then raised 100µm and left in place for another 5min to reduce upwelling.

### Ovariectomy

Mice were bilaterally ovariectomized under isoflurane anesthesia 2d, 1wk or 6wk before optogenetic studies depending on design, detailed below. At the time of surgery or five weeks after, some mice also received a Silastic implant containing either 0.625µg 17ß-estradiol in sesame oil vehicle (OVX+E) or just vehicle (OVX) subcutaneously in the scapular region; the investigators were blind to treatment. Bupivacaine (0.25%, APP Pharmaceuticals) was applied to incision areas as an analgesic for these procedures.

### Experimental design

Optogenetic stimulation and electrophysiologic recordings in acutely-prepared mouse brain slices were used to study the effects of activation of AVPV kisspeptin neurons on synaptic currents in GnRH neurons and/or GnRH neuron firing rate. Stereotaxic injection of AAV5-Syn-FLEX-rc[ChrimsonR-tdTomato]) was used to drive ChrimsonR expression specifically in kisspeptin neurons in the AVPV. First, specificity of transfection was assessed via immunofluorescence in Kiss-Cre/Kiss-GFP mice. Second, electrophysiologic recordings of transfected kisspeptin neurons, identified by co-expression of GFP and tdTomato, were done during wide-field photostimulation to determine photostimulation efficacy. Third, Kiss-Cre/GnRH-GFP mice were similarly prepared and four experiments were done to measure responses of GFP-identified GnRH neurons to optogenetic kisspeptin neuron activation. In Experiments 1 and 2, mice were OVX and given vehicle or E implants, then brain slices were prepared two days later in either the morning (AM) or late afternoon (PM). In Experiment 1, whole-cell current-clamp recordings of GnRH neurons were used to measure photostimulation-induced changes in firing rate, then voltage-clamp recordings were done in the same cells to assess photostimulation-evoked synaptic transmission. In Experiment 2, extracellular recordings were used instead of current-clamp to measure firing rate, and whole-cell voltage-clamp recordings of synaptic currents were performed on separate cells. In addition, some cells were subjected to sequential extracellular followed by whole-cell voltage-clamp recordings. In Experiment 3, this sequential extracellular then whole-cell voltage-clamp recording paradigm was used in brain slices prepared from diestrous, proestrous, or 1wk post-OVX mice in the late afternoon. In Experiment 4, mice were injected with virus, OVX 1wk later, then treated with E or vehicle implants 5wks later. One week after steroid manipulation, brain slices were prepared in the late afternoon and extracellular recordings were used to measure photostimulation-induced changes in GnRH neuron firing rate. A list of the number of mice and the number of GnRH neurons for each recording type is provided in Table 1.

**Table 1.** Number of GnRH neurons and mice per group by recording type for each Experiment. CC, current-clamp; VC 20, voltage-clamp with 20mM Cl^−^; VC, voltage-clamp with 140mM Cl^−^; EC extracellular. Grey identifies recording approaches.

| Experiment | group | cells | mice | recording |
| --- | --- | --- | --- | --- |
| 1 | OVX AM | 5 | 1 | CC/VC 20 |
|  | OVX PM | 13 | 3 | CC/VC 20 |
|  | OVX+E AM | 5 | 1 | CC/VC 20 |
|  | OVX+E PM | 1 | 1 | CC/VC 20 |
|  | OVX+E AM | 1 | 1 | CC |
| 2 | OVX AM | 4 | 2 | VC |
|  | OVX PM | 20 | 4 | VC |
|  | OVX+E PM | 29 | 5 | VC |
|  | OVX AM | 13 | 4 | EC |
|  | OVX PM | 9 | 4 | EC |
|  | OVX+E AM | 8 | 5 | EC |
|  | OVX+E PM | 12 | 4 | EC |
|  | OVX AM | 3 | 3 | EC/VC |
|  | OVX PM | 5 | 2 | EC/VC |
|  | OVX+E AM | 6 | 3 | EC/VC |
|  | OVX+E PM | 7 | 2 | EC/VC |
| 3 | diestrus | 33 | 11 | EC/VC |
|  | diestrus | 4 | 3 | EC |
|  | proestrus | 30 | 11 | EC/VC |
|  | proestrus | 2 | 2 | EC |
|  | OVX 1 wk | 27 | 9 | EC/VC |
|  | di no opsin | 16 | 5 | EC |
|  | di repeat stim | 2 | 2 | EC |
| 4 | OVX 6wk | 9 | 6 | EC |
|  | OVX+E 6wk | 10 | 6 | EC |

### Immunofluorescence

Kiss-Cre/Kiss-hrGFP mice injected with AAV5-Syn-FLEX-rc[ChrimsonR-tdTomato] (n=2) were anesthetized with isoflurane and transcardially perfused with PBS (15–20 mL) then 10% neutral-buffered formalin (∼50mL). Brains were placed into 10% formalin overnight, followed by 15% sucrose for ≥24h then 30% sucrose for ≥24h for cryoprotection. Brains were then rapidly frozen in optical cutting temperature compound and sections (30µm, four series) were cut on a cryostat (Leica CM3050S) and stored at −20°C in antifreeze solution (25% ethylene glycol, 25% glycerol in PBS). Sections were washed with 0.1M phosphate-buffered saline containing 0.1% TritonX-100 (PBST, pH 7.4), then treated with 0.1% sodium borohydride (NaBH4) in PBST for 10min. After this, slices were washed with PBST 3 times for 5min. Sections were placed in blocking solution (PBST with 4% normal goat serum, Jackson Immunoresearch) for 1h at room temperature (21-23C), then incubated with rabbit anti-hrGFP (Agilent, 1:5000) and chicken anti-mCherry (Abcam, 1:2000) in blocking solution overnight at 4°C. Sections were then washed with PBST and incubated in PBST containing Alexa Fluor 594-conjugated goat anti-chicken and Alexa Fluor 488-conjugated goat anti-rabbit antibodies (1:500, Jackson Immunoresearch) for 90min at room temperature on a shaker at 60RPM. Sections were then washed with PBS and mounted on Superfrost+ slides (Thermofisher) for imaging with a fluorescent microscope. The number of immunoreactive GFP-only, tdTomato-only, and GFP/tdTomato cells were counted in the AVPV and arcuate nuclei.

### Polymerase chain reaction (PCR)

PCR was used to examine kisspeptin mRNA levels in mice from the long-term OVX studies. Adult female mice underwent bilateral OVX. Five weeks later, mice received either estradiol (n=4) or blank (n=6) implants. One week later, a tissue block including the AVPV was collected in the morning to measure kisspeptin mRNA. Tissue was immediately homogenized in RLT buffer (QIAGEN) containing 2-mercaptoethanol (1%v/v, Sigma), snap frozen, and stored at −80°C. Total RNA was isolated with on-column DNasing (RNeasy; Qiagen, Germantown, MD) from lysates. RNA isolated from the tissue (500 ng) and a standard curve of mouse POA RNA 1250ng-10pg (1:5 dilution) were reverse transcribed with Superscript Vilo IV (Invitrogen, Waltham, MA). POA and Arc cDNA (8 ng/µL final concentration for samples) were assayed in duplicate for: *Kiss1, Actb,* and *Rps29* messenger RNAs (mRNAs) by hydrolysis probe-based quantitative PCR chemistry (TaqMan, *Actb,* and *Rps29*) or SYBR Green (*Kiss1*). All primers and probes were purchased from Integrated DNA Technologies (Coralville, IA). PCR was conducted using Applied Biosystems Gene Expression Mastermix (ThermoFisher). Normalized relative expression of transcripts was determined by the ΔΔCt method (Bustin, 2002), the average of *Actb* and *Rps29* expression were used for normalization.

### Brain slice preparation

All solutions were bubbled with 95% O_2_/5% CO_2_ for at least 15min before exposure to tissue. For Experiments 1 and 2 examining time of day, brain slices were prepared at either Zeitgeber time (ZT) 6 (“AM”) or ZT12 (“PM” expected LH surge peak (Christian et al., 2005)). For Experiment 3, brain slices were prepared at ZT12. For Experiment 4, brain slices were prepared at ZT 6. Blood was collected for serum LH measurements from all mice and the brain rapidly removed and placed in ice-cold sucrose saline solution containing in mM: 250 sucrose, 3.5 KCl, 25 NaHCO_3_, 10 D-glucose, 1.25 Na_2_HPO_4_, 1.2 MgSO_4_, and 3.8 MgCl_2_, and adjusted to pH 7.6 and 345mOsm. Sagittal (Experiment 1-2) or coronal (Experiments 2-4) slices (300μm) were cut with a VT1200S vibrating microtome (Leica Biosystems). Slices were incubated for 30min at room temperature in a 1:1 mixture of sucrose saline and artificial CSF (ACSF) containing in mM 135 NaCl, 3.5 KCl, 26 NaHCO_3_, 10 D-glucose, 1.25 Na_2_HPO_4_, 1.2 MgSO_4_, and 2.5 CaCl_2_, and adjusted to pH 7.4 and 305mOsm room temperature (∼21–23°C). Slices were then transferred to 100% ACSF for an additional 30–180min at room temperature before recording.

### Electrophysiology

Slices were placed into a chamber and perfused (3ml/min) with ACSF kept at 31°C with an inline heating unit (Warner Instruments). All recordings were performed with an EPC-10USB patch-clamp amplifier controlled with PatchMaster software (HEKA Elektronik). Recording pipettes were pulled from borosilicate capillary glass using a Flaming/Brown P-97 puller (Sutter Instruments) to obtain pipettes with a resistance of 2.5-3.5MΩ, which were wrapped with Parafilm^®^ to reduce capacitive transients. At the end of a recording session, the pipette was left in place and brightfield (differential interference contrast) and fluorescent (GFP, tdTomato) live images were taken using 4x and 40x objectives to document the location of the recorded neuron in the brain slice and to visualize cellular morphology and proximity to tdTomato-expressing fibers.

### Whole-cell recordings

Whole-cell recordings of GFP-identified GnRH neurons were made with pipettes filled with one of two solutions. For Experiment 1, current- and voltage-clamp recordings were made with a pipette solution containing a physiologic chloride concentration for GnRH neurons (Defazio 2002). This solution contained (in mM): 120 K gluconate, 20 KCl, 10 HEPES, 5 EGTA, 0.1 CaCl_2_, 4MgATP, and 0.4NaGTP at 305 mOsm, pH to 7.2 with NaOH. For Experiment 2, voltage-clamp recordings were made with a pipette solution with a near isotonic chloride concentration to increase the driving force for GABAergic synaptic currents at a holding potential of -65mV. This solution contained (in mM): 140 KCl, 10 HEPES, 5 EGTA, 0.1 CaCl_2_, 4 MgATP, and 0.4 NaGTP. All potentials reported were corrected on-line for a calculated liquid-junction potential of -14.5mV (20 mM) or -4.9mV (140 mM) (Barry, 1994). After achieving the on-cell configuration with seal resistance >2.0GΩ, fast capacitive transients were minimized and measured, and the whole-cell configuration was achieved by rupturing the cell membrane with brief suction. Passive membrane properties to assess recording quality were calculated from the averaged response after on-cell capacitive current subtraction) to sixteen -5mV, 20ms test pulses from a holding potential of -65mV performed in voltage-clamp mode immediately before and after each protocol. The following quality control criteria were required for analysis: series resistance (Rs) <25MΩ, input resistance (Rinput) >500MΩ, stable capacitance (Cm) between 8 and 30pF, holding current (Ihold) between -60 and 10pA.

### Extracellular recordings

Targeted extracellular recordings were made to obtain firing patterns of GnRH-GFP or Kiss-Cre/Kiss1-hrGFP neurons (Nunemaker et al., 2003). This method maintains the cell’s cytosolic milieu and has minimal impact on the spontaneous firing rate of neurons (Alcami et al., 2012). Pipettes were filled with a solution containing (in mM) 150 NaCl, 10 HEPES, 10 D-glucose, 2.5 CaCl2, 1.3 MgCl2, and 3.5 KCl, at pH 7.4 and 310mOsm. GFP-expressing neurons were identified by brief (<2s) illumination at 470nm. Low-resistance (< 50MΩ) seals were formed between the pipette and neuron after first exposing the pipette to slice tissue outside the area of interest in the absence of positive tip pressure. Recordings were made in voltage-clamp mode with a 0mV pipette holding potential. Signals were filtered at 8.5kHz and acquired at 20kHz. Seal resistance was checked frequently during the first three min of recordings to ensure a stable baseline. The offset potential was adjusted every minute to reset the baseline current to 0pA. Cells that exhibited no spikes during recording were excluded from analysis, as the recording could not be verified (n=5 total; 2 OVX PM, 1 OVX+E PM, 1 OVX 6wk, 1 OVX+E 6wk).

### Photostimulation hardware and AVPV kisspeptin neuron spike fidelity studies

A custom light-emitting diode (LED) system was coupled to the epifluorescence port of an Olympus BX51 microscope and used for widefield illumination and optical stimulation of brain slices via a 40x LUMPlanFl/IR water-immersion objective. For GFP excitation, a 470nm (“blue”) LED was coupled with a 470nm (40nm FWHM) filter. For excitation of tdTomato, a neutral white (4100K) LED was coupled with a 542nm (20nm FWHM) filter. For excitation of ChrimsonR, the white LED was coupled with a 620nm (58nm FWHM) filter. LEDs were controlled using the analog voltage output of the EPC-10USB connected to an externally-dimmable direct-current driver (Luxeonstar). Peak optical power was measured using Thorlabs PM101/S170C power meter to be 15.5mW/mm^2^ when using the 620nm filter by placing distilled water on the sensor surface and lowering the 40x objective into the water to the typical experimental working distance relative to the sensor surface. This power was used for all stimulations.

To determine the parameter values effective for optogenetic stimulation of AVPV kisspeptin neurons, targeted extracellular recordings were made of GFP/tdTomato-coexpressing cells (n=12) in slices from ovary-intact Kiss1-GFP/Kiss1-Cre mice 3-8wks after virus injection. Trains of 30 red light pulses were applied with varying light pulse duration (0.5, 1, 2ms) and frequency (1, 5, 10, 20, 40, 80Hz). The range of these parameters was chosen based on previous studies (Klapoetke et al., 2014; Qiu et al., 2016; Piet et al., 2018). Spike fidelity was defined as the number of evoked spikes divided by the number of light pulses multiplied by 100. Trains of 30 blue light pulses were also tested because 470nm light was required to identify GFP-expressing cells, possibly driving cross-activation of ChrimsonR due to partial overlap of activation spectra. Blue light pulses at power needed to visualize GFP indeed evoked spiking of ChrimsonR-expressing AVPV kisspeptin neurons. To determine if prolonged exposure to constant blue light affected responsiveness to subsequent red-light pulses, spike fidelity in response to red light was tested before and 5min after a 15s continuous exposure to blue light. This duration exceeded typical exposures used during GFP visualization, which were <2s.

### Photostimulation paradigms for GnRH neuron recordings

*Whole-cell recordings of fast synaptic input.* To identify fast synaptic input to GnRH neurons during whole-cell recordings, sixty 1ms duration red light pulse was applied at 1s intervals during voltage-clamp recordings. Pulse duration was increased 5-50ms if the neuron did not display evoked PSCs in response to 1ms pulses; extended pulses did not evoke a postsynaptic response in any of the cells tested in which 1 ms pulses failed (n=108). These recordings were made with both physiologic (Experiment 1) and isotonic pipette chloride concentrations (Experiments 2 and 3).

### Experiment 1: Whole-cell current-clamp recordings of GnRH neuron firing rate and voltage-clamp recordings of evoked PSCs (ePSCs)

Current-clamp recordings of GFP-positive neurons in sagittal brain slices from AAV-injected Kiss1-Cre/GnRH-GFP mice were performed to determine downstream effects of AVPV kisspeptin neuron photoactivation on GnRH neuron firing rate. Recordings were performed in sagittal brain slices from mice 2d post OVX or OVX+E in the AM or the PM in 25 cells from seven mice. Current was applied and adjusted over a 1-min stabilization period so that each recording began at approximately -65 mV (mean±SD: 63.3±1.3 mV). Membrane potential was then recorded continuously for 1min followed by a 10Hz (1ms pulse width), 30s photostimulus train, followed by a 5min recording of membrane potential. In non-responding cells, a longer stimulus train duration (60s, n=2 cells; 180s, n=3 cells) or increased stimulation pulse frequency (20Hz, n=4 cells) were tested and did not cause discernable changes in response (data not shown). Amplifier mode was switched to voltage-clamp and spontaneous PSCs and responses to 60 trials of 1ms red light pulses at 1s intervals recorded.

### Experiment 2 Whole-cell voltage-clamp recordings of ePSCs and extracellular recordings of firing response

Experiment 1 results had a low signal to noise ratio for PSCs. We thus changed the pipette solution to 140mM Cl^−^ to increase driving force; this precluded interpretable current-clamp recordings in the same cell. Voltage-clamp recordings were performed on GFP-positive neurons in sagittal slices from AAV-injected Kiss1-Cre/GnRH-GFP mice 2d post OVX or OVX+E. Responses to 60 trials of 1 ms red light pulses at 1s intervals were recorded.

In the same animal model, extracellular recordings were performed in sagittal or coronal slices to determine downstream effects of AVPV kisspeptin neuron photoactivation on GnRH neuron firing rate. Once stabilized, extracellular recordings of spontaneous action currents were made for a 2-min baseline. Then, three red-light stimulus trains with the following parameters were applied: 1^st^ train, 1ms pulses at 10Hz for 20s; 2nd train, replicate of first train; 3rd train, 1ms pulses at 20Hz for 30s. There were 5min between the start of these trains and recordings continued for five min after the end of the third train. No differences were noted with slice orientation,

While the changes in approach used for Experiment 2 mitigated some noted caveats in Experiment 1, we still wanted to test if the presence/absence of a stimulus-induced change in firing rate was predictive of the presence/absence of a synaptic connection, as indicated by the ability to photo-evoke PSCs. To do this, sequential recordings were performed in some cells. At the end of the extracellular recording, the pipette was carefully withdrawn and a new pipette with 140 mM Cl^−^ solution used to perform a whole-cell recording of the response to 60 1ms red light pulses at 1s intervals from the same cell.

### Experiment 3: Extracellular recordings of firing response and whole-cell voltage-clamp recordings of ePSCs in diestrous, proestrous, and 1wk post-OVX mice

Experiment 2 suggested the 20Hz, 30s train was most effective at evoking a change in firing rate, but it was both longer duration and higher frequency than the preceding 10 Hz stimulus trains. For Experiment 3, we held duration at 30s while varying the frequency of light pulses to determine the effect of frequency. Mice in diestrus, proestrus or 1wk post OVX were tested at frequencies of 5, 10, or 20Hz (30s train duration, 1ms pulse duration) in varied order with 15min intertrain intervals and 15min of stimulus-free recording before the first train and after the end of third train. Following the extracellular recordings, the pipette was carefully withdrawn and voltage-clamp recordings attempted as in sequential recordings in Experiment 2 with 140 mM Cl^−^. To test if ePSCs were monosynaptic and to determine the mediating neurotransmitter (n=5) (Petreanu et al., 2009), sequential application of CNQX (10µM, this treatment absent in one cell), tetrodotoxin TTX, 0.5µM), 4-aminopyridine (4-AP, 0.5mM), gabazine (5µM), then ACSF for washout was done. Additional cells were treated only with CNQX followed by gabazine (n=3) to test the mediating neurotransmitter. All pharmacology experiments were performed in coronal brain slices made in the PM from diestrous or proestrous mice. At the end of whole-cell recordings without pharmacological treatments, some cells were given a 20Hz, 30s stimulus train to assess PSCs during high frequency stimulation.

### Experiment 4: Extracellular recordings of firing response in 6wk post-OVX mice with or without 1wk estradiol treatment

To test if longer-term OVX, which is reported to reduce kisspeptin immunoreactivity in the AVPV (Brock and Bakker, 2013), would reduce GnRH neuron response to a 20Hz, 30s stimulus, mice received stereotaxic injections as above and were OVX 1wk later. Five wks after OVX, mice received either VEH or E implants, and recordings were performed 1wk later.

### Data analysis

For extracellular recordings, action currents (spikes) and PSCs were detected off-line using custom scripts in Igor Pro 9 (Sutter Instruments). To determine if individual neurons showed a change in firing rate in response to photostimulation trains, a within-cell permutation test, described below, was performed on data divided into 20s bins. The period of the stimulus was excluded from these tests but was examined separately for cells initiating firing or exhibiting a marked increase in firing rate within 5s of photostimulus onset compared to the 5s immediately preceding the stimulus. For current-clamp recordings (Experiment 1), only 1min of recording preceded the first stimulus and most cells were silent during this baseline period, making a permutation test ineffective for detecting changes in firing. Therefore, to calculate a change in firing rate, the firing rate during the 1min preceding the stimulus train was subtracted from the firing rate during the 5min after the stimulus train.

In Experiment 2, 2min of recording preceded the first 10 Hz stimulus; 4min represents the maximum window providing equal before- and after-stimulus periods. For the second 10Hz stimulus and for all 20Hz stimuli (Experiments 2, 3, and 4), an 8min window was used (4min before and after). For Experiment 2, there was a 1min overlap between the first 10Hz after-stimulus analysis period and the before-stimulus period of the next 10Hz stimulus train, and a 4min overlap between the after-stimulus period of the second 10Hz train and the before-stimulus period of the 20Hz train. For each stimulus train, the mean firing rate during the before-stimulus analysis period was subtracted from the after-stimulus analysis period to calculate an observed difference, and a permutation test used to assign a p value to this difference. To construct a null distribution of firing rate differences expected by chance, 20s bins during the before and after-stimulus periods were randomly permuted 10,000 times and a difference score was calculated for each permutation. The *P*-value was defined as the proportion of permutations in which the difference score equaled or exceeded the observed difference. *P*-values were adjusted for multiple comparisons using Benjamini-Hochberg correction. Cells were classified as responsive if adjusted *P*<0.05, indicating the observed change in firing rate was unlikely to have occurred by chance, the firing rate difference between before- and after-stimulus analysis periods exceeded 0.25Hz, and there was at least a 25% change between these periods. Notably, testing several variables (e.g., duration, bin size) of this analysis produced largely similar results that would not change biological interpretation.

For whole-cell recordings, PSCs were detected off-line using custom scripts in Igor Pro 9. Evoked PSC (ePSC) latency was calculated as the duration between the start of each light pulse and the peak of the PSC, with PSCs categorized as evoked if the latency was between 0.5 and 13ms. Initiating the range of evoked PSCs at 0.5ms eliminated spontaneous events that began before the pulse began but peaked during the light pulse; 13ms is the mean+2SD of the evoked PSC latency in presence of TTX+4AP, which defined monosynaptic events. To mitigate erroneous classification of cells with high spontaneous PSC rates as synaptically responsive, a synapse probability score was calculated based on spontaneous PSC frequency and the number of trials in which a PSC was detected within the accepted window. The probability of the recorded cell receiving monosynaptic transmission from the stimulated AVPV kisspeptin population was calculated based on a Poisson model,

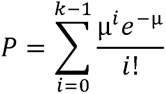

where µ is the expected number of spontaneous PSCs within the evoked window across all 60 trials, calculated as (n trials * spontaneous PSC rate in Hz * window duration in s). A PSC was considered evoked if *P*<0.05. Total charge transfer occurring during PSC recordings was calculated by integrating current over time during the 30s before, during, and after the stimulus train.

### Statistical analyses

Data were analyzed using Igor Pro 9, Prism 10 (GraphPad), and Python 3.14. Shapiro-Wilk was used to test for normality of the distributions. Data are plotted as mean ± SEM or median ± interquartile range and were compared among animal groups with statistical tests appropriate for experimental design and data distributions as indicated in the results section. The number of cells per group is indicated by n. Analysis code is provided at https://gitlab.com/um-mip/coding-project.git.

## Results

### AAV5 delivery of FLEX ChrimsonR-tdTomato constructs targets cells expressing Cre recombinase under control of the kisspeptin promoter

Brain sections from Kiss1-Cre/Kiss1-hrGFP mice injected with AAV-Syn-FLEX-rc[ChrimsonR-tdTomato] into the AVPV were immunolabeled. hrGFP immunoreactivity was observed predominantly in the AVPV and arcuate nuclei consistent with known hypothalamic kisspeptin populations and previous characterization of the Kiss1-hrGFP mouse line (Gottsch et al., 2004; Lehman et al., 2010; Cravo et al., 2013). tdTomato immunoreactivity was observed in cell bodies and fibers in the AVPV but fibers only in the arcuate with the exception of 3 of 772 cells examined. Dual immunoreactive hrGFP/tdTomato cells were 76.6 and 83.3% of hrGFP-expressing cells and <6% of cells in the AVPV expressed only tdTomato. These results indicated the AAV vector drove Cre-dependent ChrimsonR/tdTomato expression in AVPV kisspeptin neurons as has been reported (Wang et al., 2019).

### Photoactivation of AVPV kisspeptin neurons generates high-fidelity spike trains

Extracellular recordings were used to assess the ability of photostimulation of ChrimsonR to drive action currents (“spikes”) in GFP/tdTomato-expressing neurons in sagittal brain slices from AAV-injected Kiss1-Cre/Kiss1-hrGFP mice (Figure 2A). High spike fidelity (>95%) was maintained up to 20Hz, but was reduced at photostimulus frequencies ≥40Hz (Figures 2B, C). Pulse width was varied (0.5, 1, or 2ms), but had no effect within the range tested and there was no interaction of pulse width and stimulation frequency (two-way, repeated-measures ANOVA; Table 2). A pulse width of 1ms was chosen for the remainder of experiments using extracellular recordings.

**Figure 1.**
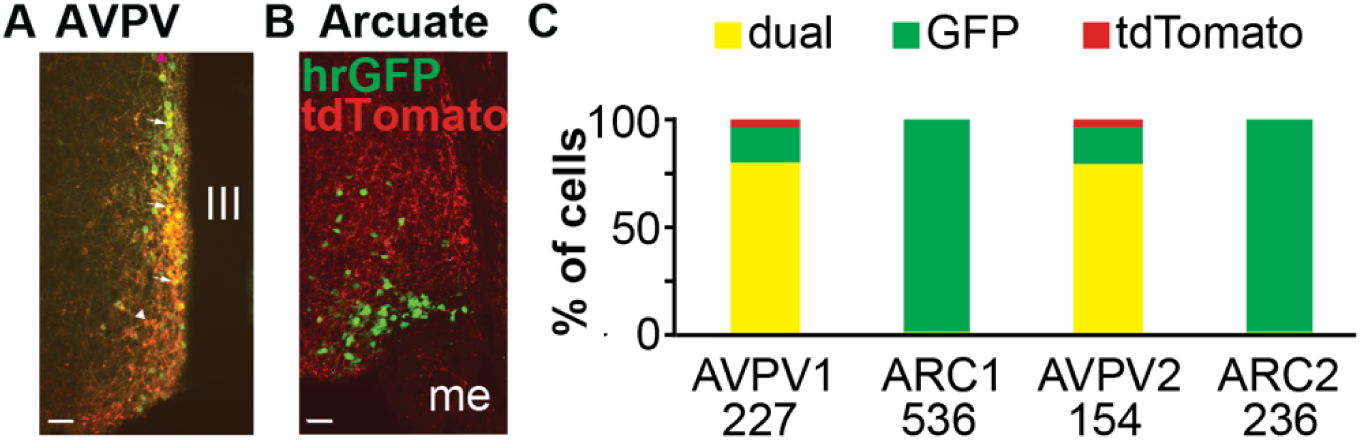
AAV5 delivery of FLEX ChrimsonR-tdTomato effectively targets kisspeptin neurons in Kiss-Cre mice. A and B, Representative dual immunofluorescent images of the AVPV (A) and arcuate (B). White arrows point to examples of colocalization, white arrowhead to a tdTomato only cell and magenta arrowhead to a GFP-only cell. C. Quantification of infected cells expressing both markers vs individual markers in both mice, number of cells per region is shown at the bottom. Scale bars are 50µm. III, third ventricle; me, median eminence.

**Figure 2.**
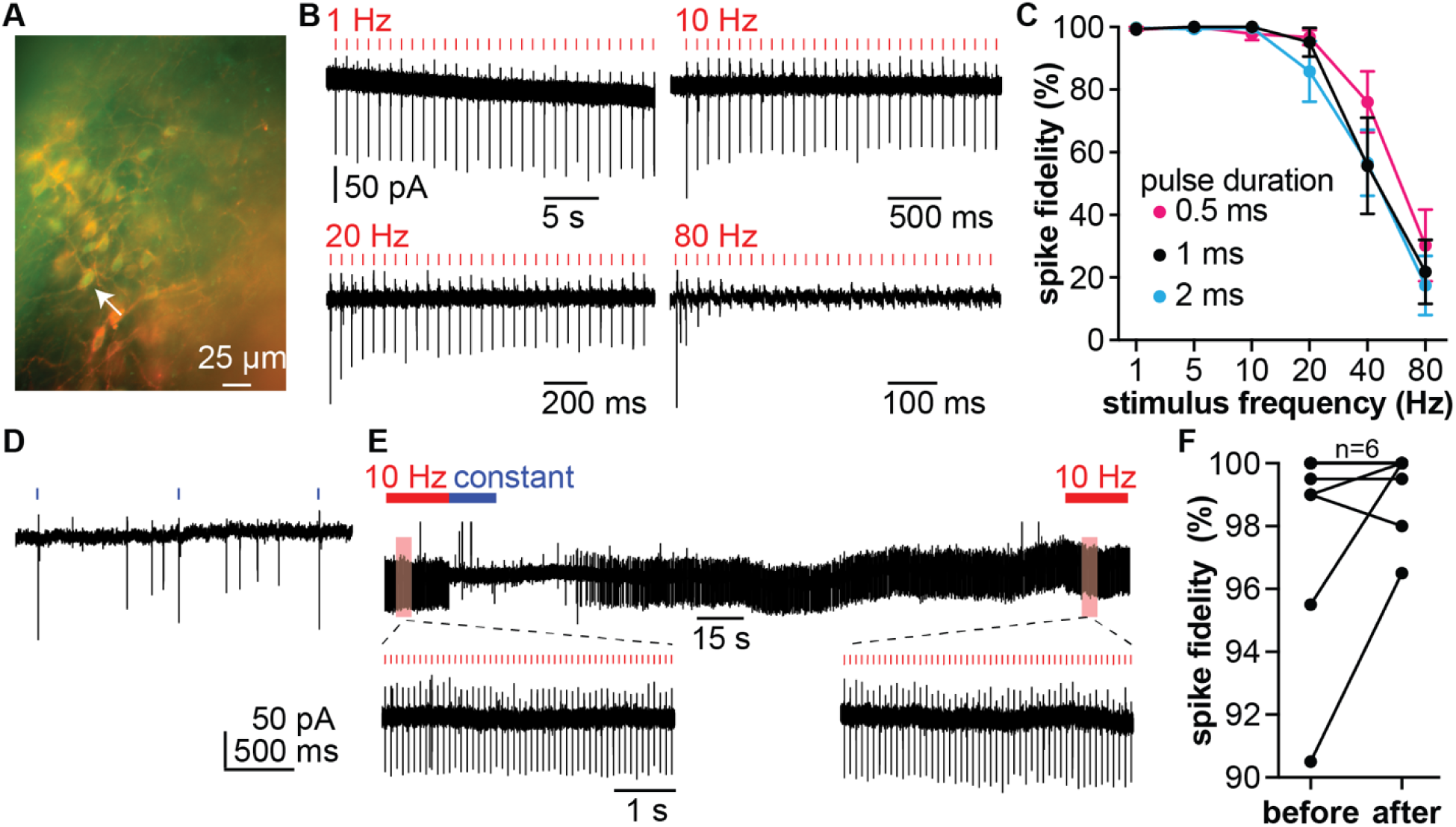
Photostimulation drives spiking in ChrimsonR-expressing AVPV kisspeptin neurons. A. Image of a 300µm sagittal brain slice from AAV-injected Kiss1-Cre/Kiss1-hrGFP mouse. Arrow indicates an example GFP/tdTomato co-expressing neuron. B. Representative extracellular recordings showing responses of an AVPV Kiss1-GFP/Kiss1-Cre neuron to thirty 1ms red-light pulses delivered at 1, 10, 20, and 80Hz. Time scale varies but 50pA scale bar applies to all traces in this figure. C. Mean±SEM spike fidelity as a function of photostimulus frequency for 0.5, 1, and 2ms red-light pulses. D. Representative extracellular recording showing response of a ChrimsonR-expressing AVPV kisspeptin neuron to three 1ms blue-light pulses at 1Hz; spontaneous spikes occurred between light pulses. E. Representative extracellular recording testing the effects of 15s constant blue light on spike fidelity. Expanded regions (red boxes in upper trace) show light-evoked spikes during each fidelity test. F. Spike fidelity before and after constant blue-light exposure. Note y-axis begins at 90%. Data for 11 neurons are shown; 6 neurons exhibited 100% fidelity before and after blue light exposure (n=6) and they are obscured by overlap.

**Table 2.**
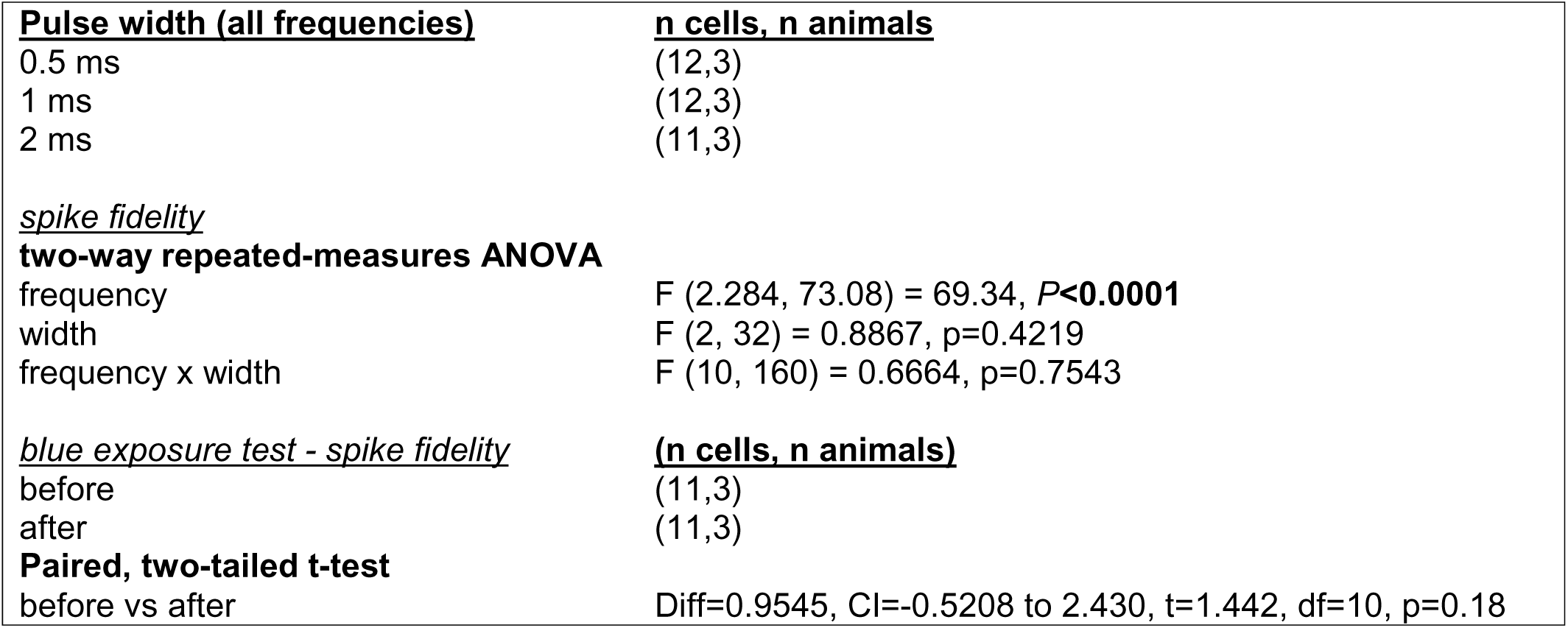
Sample sizes and statistical test results for spike fidelity and blue-exposure experiments in Figure 2. Bold text marks *P*<0.05.

The effects of exposure to blue light were also tested to determine if GFP excitation drove cross-activation of ChrimsonR. Blue light pulses applied at the power necessary to visualize GFP-expressing cells induced spikes in tdTomato/GFP co-expressing neurons (Figure 2D). To test if prolonged photoactivation by blue light affected responsiveness to subsequent photoactivation by red light, spike fidelity was examined before and after 15s constant blue light exposure (Figure 2E). Mean ±SEM spike fidelity was 98.5±3.0% before blue light exposure and 99.45±1.2% after exposure (Figure 2F). This suggested blue light (<∼2s) used for GFP visualization to identify neurons for recording was unlikely to impact the spike fidelity of subsequent red-light stimulus trains.

AAV-injected Kiss1-Cre/GnRH-GFP mice were used for remaining studies to examine AVPV kisspeptin-to-GnRH neuron communication and postsynaptic firing response of GnRH neurons in four experiments.

### Experiment 1

Whole-cell recordings were performed of GnRH neurons in sagittal brain slices to estimate the degree of functional fast synaptic connectivity between AVPV kisspeptin and GnRH neurons, to examine the GnRH neuron firing response following photoactivation of kisspeptin neurons, and to determine if either parameter is regulated by circulating estradiol and/or time of day. Mice were OVX±E and brain slices prepared two days later, either in the AM (ZT6) or at the predicted peak of the LH surge (ZT12). GnRH-GFP neurons (OVX AM n=5 cells, OVX PM n=13, OVX+E AM n=5 plus n=1 current-clamp only, OVX+E PM n=1) with apposed tdTomato/ChrimsonR positive fibers were targeted for recording (Figure 3A). For these recordings, the pipette solution contained a physiologic chloride concentration to preclude artificial excitation of neurons by GABAergic inputs during current-clamp recordings. In current-clamp mode, most cells (n=17 of 25) exhibited no spiking activity. Two cells (OVX AM and OVX+E AM, grey in Figure 3C) exhibited no spiking before or during photostimulation and <8 spikes occurred >1 min after the photostimulation terminated; these were considered non-responding cells as the activity was both minimal and not proximal to the photostimulation. The remaining six cells all exhibited higher firing rate after the stimulus than before the stimulus (Figure 3B left, mean±SEM change in firing rate 0.78±0.48Hz n=3 OVX AM, 1.16±0.59 n=2 OVX+E AM, 1.59Hz n=1 OVX PM. Repeating the stimulus train with similar parameters (n=22 cells) or increasing the stimulus train duration and/or frequency (n=9 cells) within the ranges tested did not evoke firing rate increases when the initial stimulus train did not (data not shown). After current-clamp recordings, the amplifier was switched to voltage-clamp and fast-synaptic responses to light pulses were assessed (Figure 3B, right). Only four of 24 cells tested examined exhibited light-evoked PSCs (OVX AM: 0 of 5, OVX+E AM: 2 of 5, OVX PM: 2 of 13, OVX+E PM: 0 of 1). Interestingly, these were not always cells that exhibited an increase in firing rate. The change in firing rate from before to after stimulus and synapse/no-synapse classification is shown for each cell in Figure 3C. Statistical power was inadequate for testing interactions across hormone, time of day, frequency, and duration factors.

**Figure 3.**
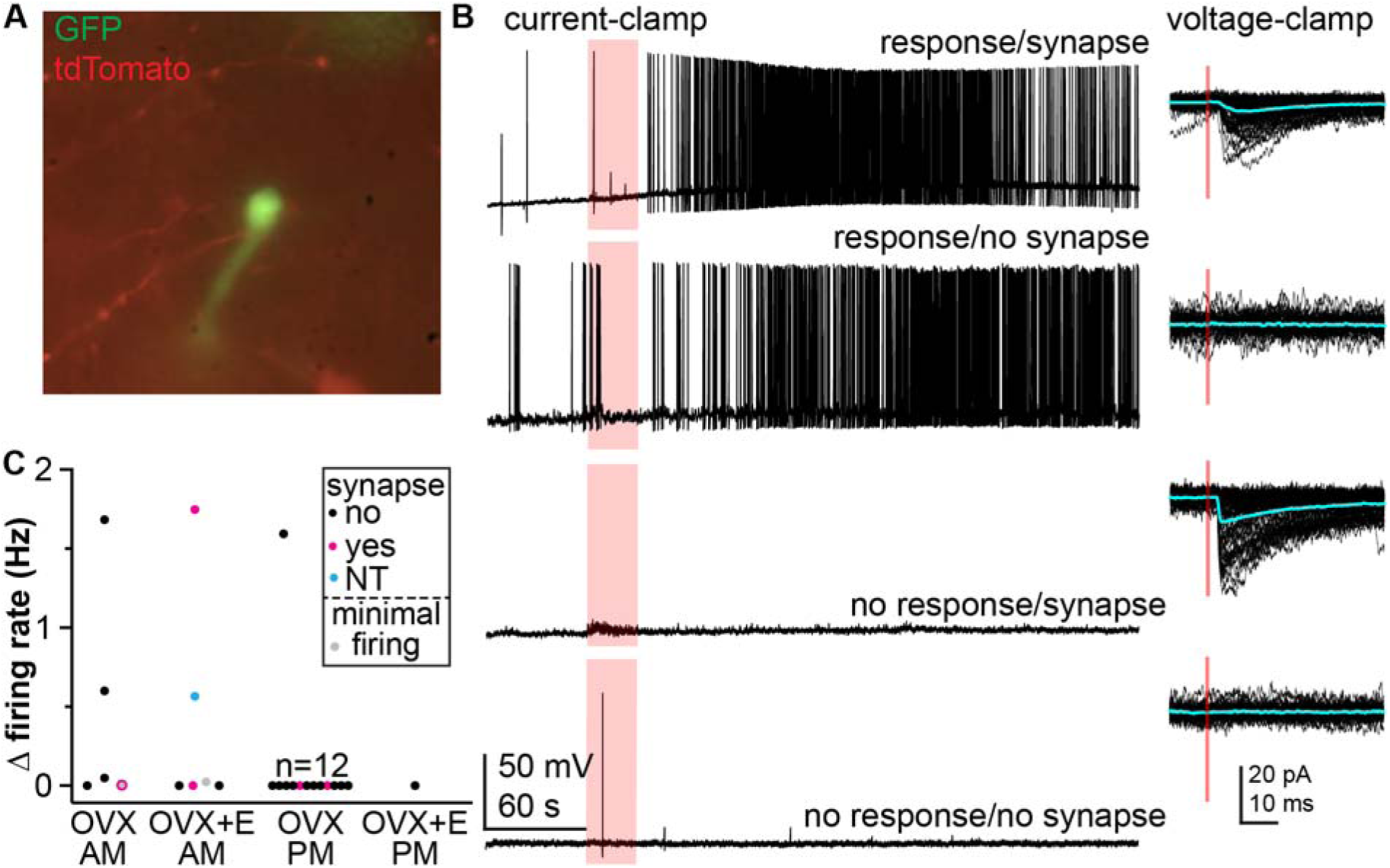
Photoactivation of AVPV Kiss1-Cre neurons during whole-cell recordings of GnRH-GFP neurons in the daily LH surge model. A. Image of a GFP-identified GnRH neuron targeted for recording in a sagittal brain slice; GnRH-GFP neurons with proximal ChrimsonR-tdTomato positive fibers were targeted for recordings. B. Example current-clamp recordings of four GnRH neurons classified by firing response to 10Hz/30s stimulus (left) and synapse during voltage-clamp recording (right) as shown above each trace. On the right, responses to individual 1ms light pulses (red) are overlaid with the mean shown in cyan. C. Change in firing rate for each cell recorded using this paradigm are color coded for presence/absence of evoked postsynaptic currents, the two cells with minimal firing post stimulus are in grey. NT, not tested.

The observation that a firing response was not always associated with detection of a synapse and that detection of a synapse did not predict a firing response was intriguing and motivated us to increase the rigor of our design to determine if this was an artifact of our experimental conditions. Specifically, the low percentage of GnRH neurons exhibiting either a change in firing rate or evoked PSCs raised three primary concerns about our approach of using whole-cell recordings with physiologic chloride concentration. First, the signal to noise ratio for PSCs was lower due to a low driving force on chloride, making it possible that evoked PSCs were missed. Second, the whole-cell configuration dialyzes the cytosol, which may dilute secondary messengers necessary for neuromodulator signaling. Third, regulating baseline membrane potential toward -65mV by injecting direct current prior to the start of each current clamp recording could interfere with spontaneous firing rates.

### Experiment 2

To address the above concerns, we performed extracellular recordings of GnRH-GFP neurons while photoactivating AVPV kisspeptin neurons in sagittal or coronal slices prepared in the morning (ZT4) or afternoon (ZT12). Following a 2min recording of baseline spike rate, three photostimulus trains were applied with 5min intertrain intervals: 10Hz1 (20s), 10Hz2 (20s), and 20Hz (30s); recordings continued for 5min after the last stimulus train (Figure 4A). Spike rates were calculated in 20s bins (Figure 4A). There was no effect of estradiol treatment or time of day on GnRH neuron firing rate during the prestimulus baseline (Figure 4B, mixed effects repeated measures three-way ANOVA all p>0.15; Table 3). Consolidating data to test for effects of either estradiol or time of day via two-way ANOVA also revealed no differences (not shown, all p>0.16). A permutation test was used to identify after-stimulus firing patterns that represented a significant increase from the before-stimulus firing pattern. Eighteen of the 57 stimulus trains tested induced an increase in firing rate. All but five of these increases were following 20Hz stimulation. Four of the 18 responses were notably late, beginning >2min after stimulus onset (3 OVX AM, 1 OVX+E AM). Not surprisingly given the low rate of response, there was no effect of time of day or estradiol on the change in firing rate from before to after for any of the three stimuli (Figure 4C, two-way ANOVA, Table 4).

**Figure 4.**
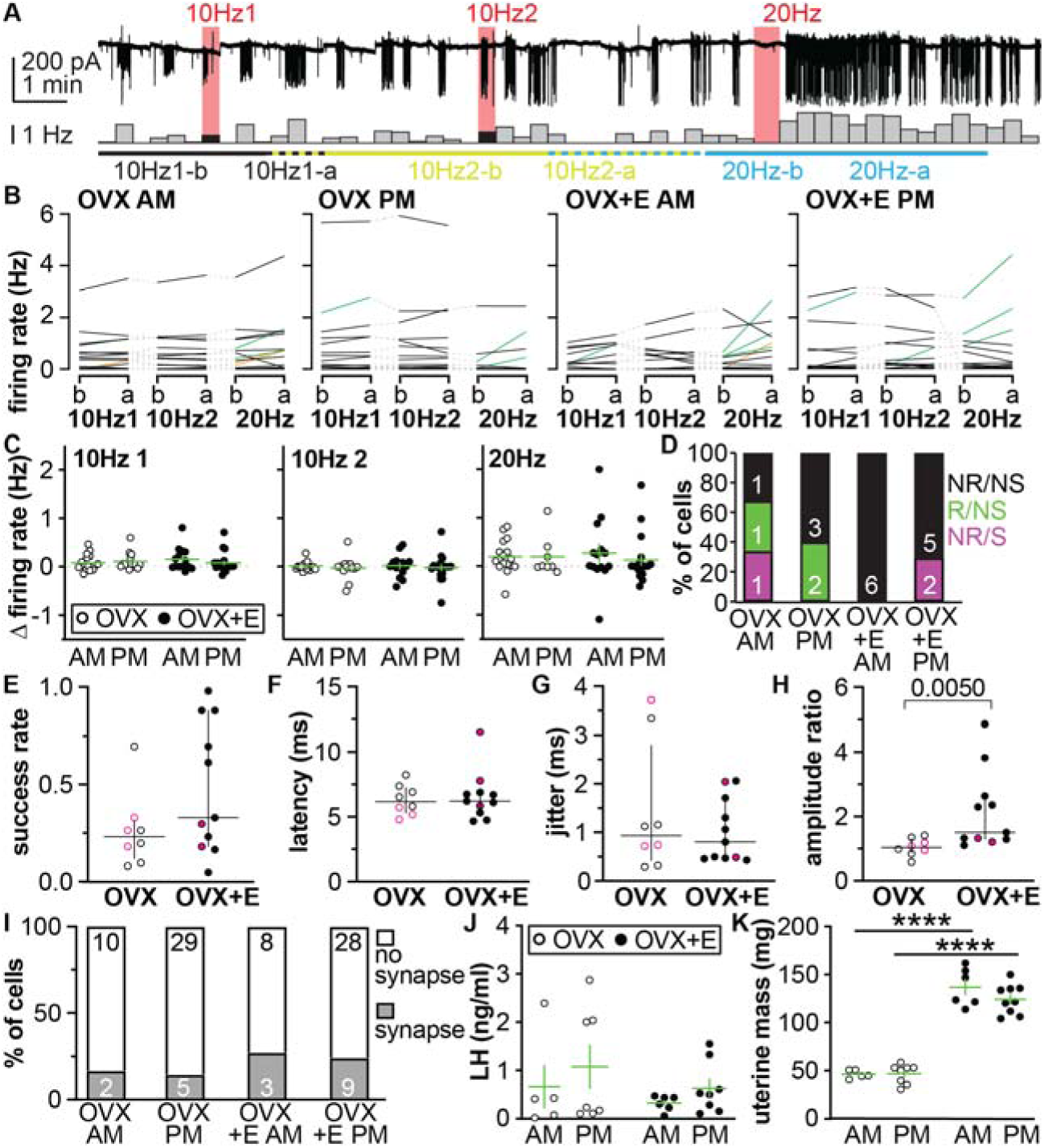
Effect of estradiol and time of day on response of GnRH neurons to AVPV kisspeptin neuron stimulation. A. Example recording trace (top) and mean firing rate in 20s bins (bottom); stimulus bins were 20s for 10Hz and 30s for 20Hz (black, excluded from analyses). Colored line below shows bins used for analysis of firing rate before (b) and after (a) each stimulus: black 2 min either side of 10Hz stimulus 1, yellow-green 4 min either side of 10Hz stimulus 2, cyan 4 min either side of 20Hz. Dashes indicate overlap of analysis periods. B. Individual values for mean firing rate before and after each stimulus; grey dashed connect the same cell. C. Change in firing rate with each stimulus. D. % of cells with sequential extracellular and whole-cell recordings by response and synapse characteristics; R response, NR no response, S synapse, NS no synapse. Number of cells indicated in bars. E-H. Individual values and median ± IQR evoked synapse success rate (E), latency (F), jitter (G) and amplitude ration (H); magenta coloring shows cells with 20mM chloride. I. % of cells for all recordings in this animal model tested for synapses that exhibited them, numbers of cells indicated in bars. J, K, Individual values and mean±SEM serum LH (J) and uterine mass (K).

**Table 3.** Three-way mixed-effects model analysis of baseline firing rate in the daily surge model.

| **Source of Variation** | **F (DFn, DFd)** | **P value** |
| --- | --- | --- |
| **stimulus** | $F(1.08, 60.47) = 2.03$ | 0.1588 |
| **time of day** | $F(1, 59) = 0.84$ | 0.3625 |
| **estradiol** | $F(1, 59) = 2.08$ | 0.1544 |
| **stim x time of day** | $F(1.08, 60.47) = 0.72$ | 0.4095 |
| **stim x estradiol** | $F(1.08, 60.47) = 0.24$ | 0.6414 |
| **time x estradiol** | $F(1, 59) = 0.40$ | 0.5316 |
| **stim x time of day x estradiol** | $F(1.08, 60.47) = 0.21$ | 0.6699 |

**Table 4.** Two-way ANOVA analysis of change in firing rate in the daily surge model.

| **Frequency** | **Source of Variation** | **F (DFn, DFd)** | **P value** |
| --- | --- | --- | --- |
| **10 Hz (1)** | time of day | $F(1, 59) = 0.24$ | 0.6274 |
| | estradiol | $F(1, 59) = 0.19$ | 0.6633 |
| | time of day x estradiol | $F(1, 59) = 1.05$ | 0.3106 |
| **10 Hz (2)** | time of day | $F(1, 59) = 0.40$ | 0.5291 |
| | estradiol | $F(1, 59) = 0.01$ | 0.9171 |
| | time of day x estradiol | $F(1, 59) < 0.01$ | 0.9959 |
| **20 Hz** | time of day | $F(1, 53) = 0.25$ | 0.6202 |
| | estradiol | $F(1, 53) < 0.01$ | 0.9905 |
| | time of day x estradiol | $F(1, 53) = 0.25$ | 0.6211 |

To obtain synaptic connectivity information, which is not possible with extracellular recordings, the same photostimulation paradigm for synapse investigation as Experiment 1 was paired with whole-cell voltage-clamp recordings using an isotonic pipette chloride solution to increase driving force on GABA_A_ receptor-mediated currents at -65mV. A total of 53 cells (OVX AM n=4, OVX PM n=20, OVX+E AM n=0, OVX+E PM n=29) were recorded; data collection was intentionally PM-biased to examine effects of estradiol in the PM, when it induces positive feedback(Christian et al., 2005). We remained interested in obtaining both types of data from the same GnRH neuron. We thus also made extracellular recordings followed by whole-cell voltage-clamp recordings of the same cell (OVX AM n=3 cells, OVX PM n=5, OVX+E AM n=6, OVX+E PM n=7). PSCs were considered evoked if they peaked from 0.5-13ms of light pulse onset (range empirically determined from experiments described below). An AVPV kisspeptin neuron was considered to synapse with a GnRH neuron if PSCS were evoked at a higher rate than would be expected due to random chance based on the spontaneous PSC rate of that cell. As in Experiment 1, the presence of a synapse was not predictive of a firing rate response (Figure 4D).

We combined cells from Experiment 1 and 2 for synapse analysis as the same animal models and synapse detection protocols were used. The percentage of GnRH neurons with detected synapses was similar between physiologic (Experiment 1) and isotonic chloride (Experiment 2) pipette solutions (5 of 24 cells [20.8%] vs 14 of 74 cells [18.9%]). The ratio of the amplitude of evoked to spontaneous currents was used to correct for driving force differences across pipette solutions. Because of the low incidence of synapses and the low number of cells examined in the AM, cells were grouped by OVX vs OVX+E for analysis of the properties of evoked currents. Estradiol did not affect the proportion of cells exhibiting synapses (X^2^ 0.7496, df=1, p=0.3866). Estradiol also had no effect on the success rate, latency or jitter of the currents (Figure 4 E-G, all p>0.5 Mann Whitney U test, Table 5). Estradiol did, however, increase the amplitude ratio (Figure 4H, p=0.0273 Mann Whitney U test, Table 5). The percent and number of cells from the four groups with detected synapses are shown in Figure 4I. There was no effect of time of day (X^2^ 0.5681, df=3, p=0.4510), or estradiol (X^2^ 3.285, df=3, p=0.3496) on number of cells with synapses detected.

**Table 5.** Mann-Whitney U test of evoked PSC characteristics in cells from OVX vs OVX+E mice in the daily surge model.

| **Measurement** | **OVX Median (IQR)**<br>**(n=8)** | **OVX+E Median (IQR)**<br>**(n=11)** | **Mann-Whitney U** | **P value** |
| --- | --- | --- | --- | --- |
| **success rate (%)** | 23.33 (19.59) | 33.33 (70.00) | 28.50 | 0.2124 |
| **latency (ms)** | 6.18 (1.95) | 6.22 (1.57) | 44.00 | > 0.9999 |
| **jitter (ms)** | 0.94 (2.37) | 0.81 (1.24) | 42.00 | 0.9039 |
| **amplitude ratio** | 1.04 (0.44) | 1.50 (1.35) | 11.00 | 0.0046** |

Notably, serum LH measured in blood collected from OVX+E mice recorded in the afternoon was similar to that of OVX+E mice recorded in the morning and were indeed lower in estradiol-treated mice than OVX mice, suggesting estradiol implants had not induced positive feedback at the time brain slices were prepared in the PM as is typically observed in this model (Figure 4J, two-way ANOVA, Sidak, Table 6). Uterine mass was greater in OVX+E than OVX mice regardless of time of day, as expected from estradiol exposure (Figure 4K, two-way ANOVA, Sidak, Table 6). These data are thus best interpreted in the context of estradiol treatment rather than a particular feedback state.

**Table 6.**
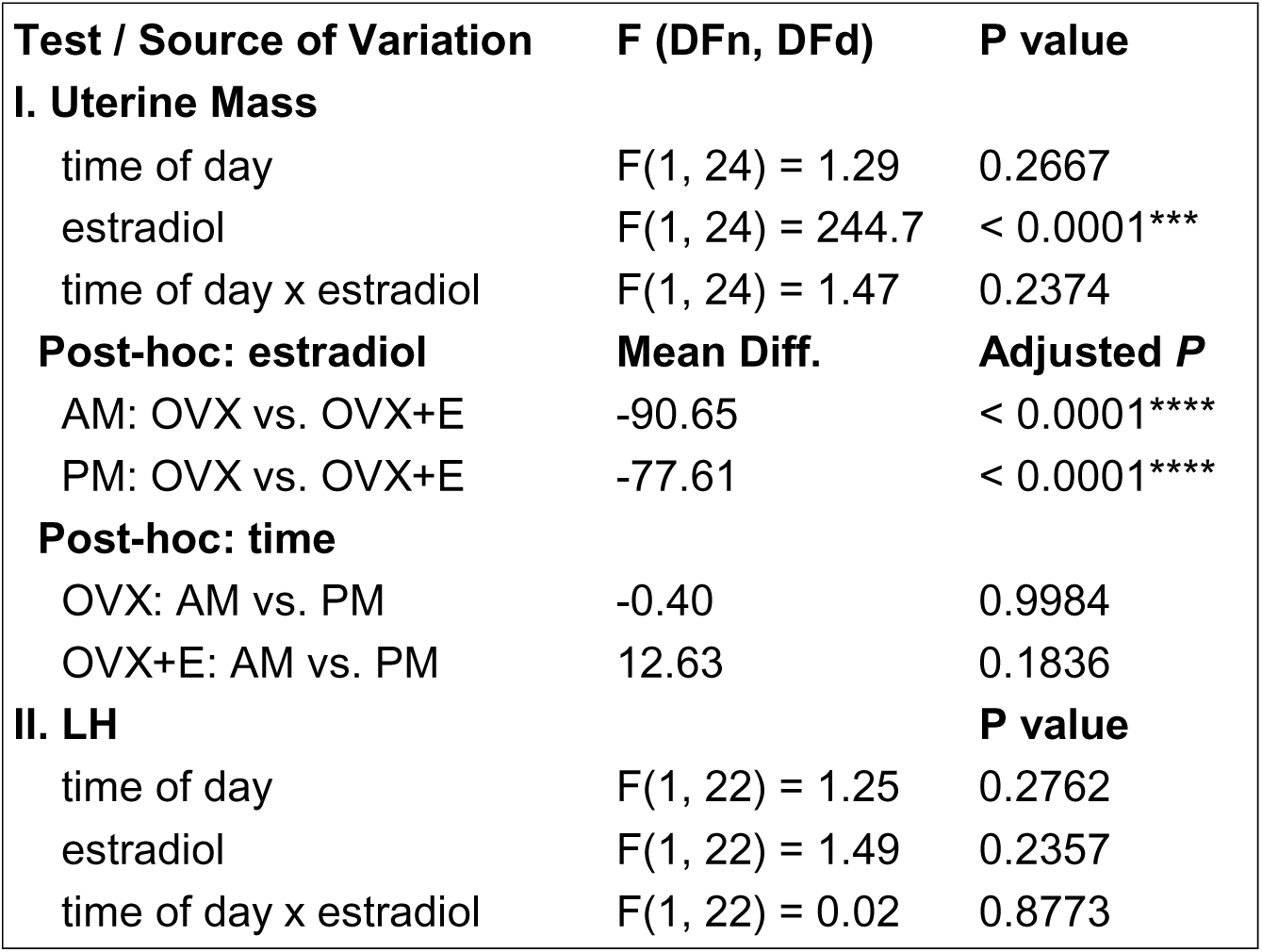
Two-way ANOVA analysis of uterine mass and LH in the daily surge model.

### Response of GnRH neurons to activation of AVPV kisspeptin neurons in brain slices from diestrous, proestrous, and ovariectomized mice

The lack of a difference in serum LH between OVX+E mice recorded in the AM vs PM suggested either failure of the estradiol treatment to induce positive feedback in the PM or preparation of brain slices before surge onset. To assess differences between conditions of estradiol negative vs positive feedback we thus switched to using ovary-intact mice, which typically have more robust LH surges(Silveira et al., 2017). We prepared brain slices in the afternoon (ZT12) from diestrous and proestrous mice as models of negative and positive steroid feedback, respectively, and from mice OVX one week before slice preparation to model the open feedback loop condition. We also examined diestrous mice with no opsin injection as negative controls. Serum LH was higher in proestrous and OVX mice than diestrous mice (Figure 5A, Kruskal-Wallis/Dunns, Table 7). Uterine mass was higher in proestrous than diestrous mice, and higher in diestrous than OVX mice (Figure 5B, Kruskal-Wallis/Dunns, Table 7). Together these measurements indicated increased estradiol exposure and induction of positive feedback in proestrous mice.

**Figure 5.**
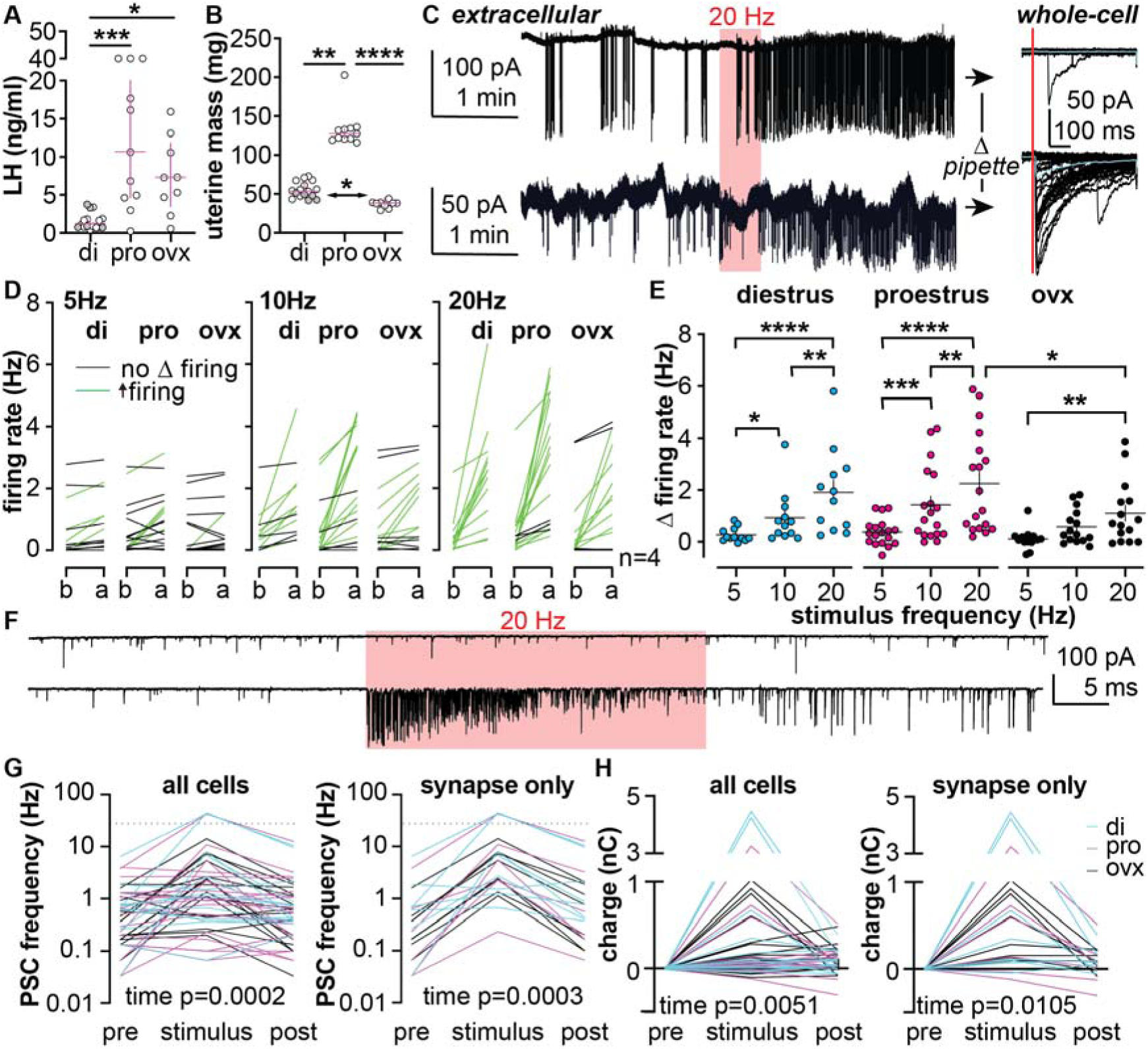
Photoactivation of AVPV Kiss1-Cre neurons in diestrous, proestrous, and 1-wk OVX mice; analysis of cells with 5, 10 and 20Hz photostimuli. A, B. Individual values and median±interquartile range for serum LH (A) and uterine mass (B). Grey symbols in the diestrous group indicate no-opsin mice. C. Left, example extracellular recordings of two GnRH neurons with photostimulus trains shown in red. Right, whole-cell voltage-clamp recordings of the same neurons; the top cell has no evoked PSC (at red line), the bottom cell has evoked PSCs. Cyan line is average of all traces shown. D. Individual values for mean firing rate before (b) and after (a) each stimulus; green indicates responding cells, black are non-responding cells. Group and stimulus rate are indicated at top. E. Individual values and mean±SEM change in firing rate (Δ). F. Representative traces of whole-cell recordings for 30sec pre stimulus (pre), during stimulus, and post stimulus (post). G. Individual values from all cells (left) and cells with detected synapses (right) of PSC frequency changes. H. Individual values from all cells (left) and cells with detected synapses (right) of changes in charge G. * *P*<0.05, ** *P*<0.01, *** *P*<0.001, **** *P*<0.0001 two-way repeated-measures ANOVA/Holm-Sidak. di, diestrus; pro, proestrus; ovx, ovariectomized.

**Table 7.** Analysis of uterine mass and LH in diestrous, proestrous, and OVX (1wk) mice.

| Test / Source of Variation | Statistic (H or Z) | df | Adjusted P value |
| --- | --- | --- | --- |
| **I. Uterine Mass** |  |  |  |
| **Kruskal-Wallis Test** | **29.75 (H)** | **2** | **< 0.0001\*\*\*\*** |
| diestrus vs. proestrus | 3.317 (Z) | — | 0.0027** |
| diestrus vs. OVX (1wk) | 2.729 (Z) | — | 0.0190* |
| proestrus vs. OVX (1wk) | 5.421 (Z) | — | < 0.0001**** |
| **II. Plasma LH** |  |  |  |
| **Kruskal-Wallis Test** | **15.74 (H)** | **2** | **0.0004\*\*\*** |
| diestrus vs. proestrus | 3.681 (Z) | — | 0.0007*** |
| diestrus vs. OVX (1wk) | 2.777 (Z) | — | 0.0165* |
| proestrus vs. OVX (1wk) | 0.634 (Z) | — | > 0.9999 |

In additional to changing the animal model, we evolved the photostimulation approach. We hypothesized hormone feedback state modulates the stimulus-response sensitivity and tested this by applying 5, 10, and 20 Hz photostimulus trains (30s) with 15min intertrain intervals. We further hypothesized the presence/absence of a synaptic connection between the AVPV kisspeptin neuron population and the recorded GnRH neuron influences the GnRH neuron’s firing rate response. To test this, we used the sequential recording approach used in Experiment 2 performing extracellular recordings followed by whole-cell recording of the same neuron (Figure 5C). Spike rate data from extracellular recordings were analyzed similar to Experiment 2. Increasing stimulus frequency increased the proportion of cells exhibiting an increased firing rate (Figure 5D, E, two-way ANOVA, Table 8). In cells from diestrous mice, for example, the proportion of cells responding to 5Hz was 4 of 13, whereas all 13 cells responded to 20Hz. The effect of increasing stimulus frequency was milder in cells from the open loop OVX mice and the change in firing rate was greater in response to 20Hz stimuli in cells from proestrous than from OVX mice (Figure 5E, two-way ANOVA, Table 8).

**Table 8.**
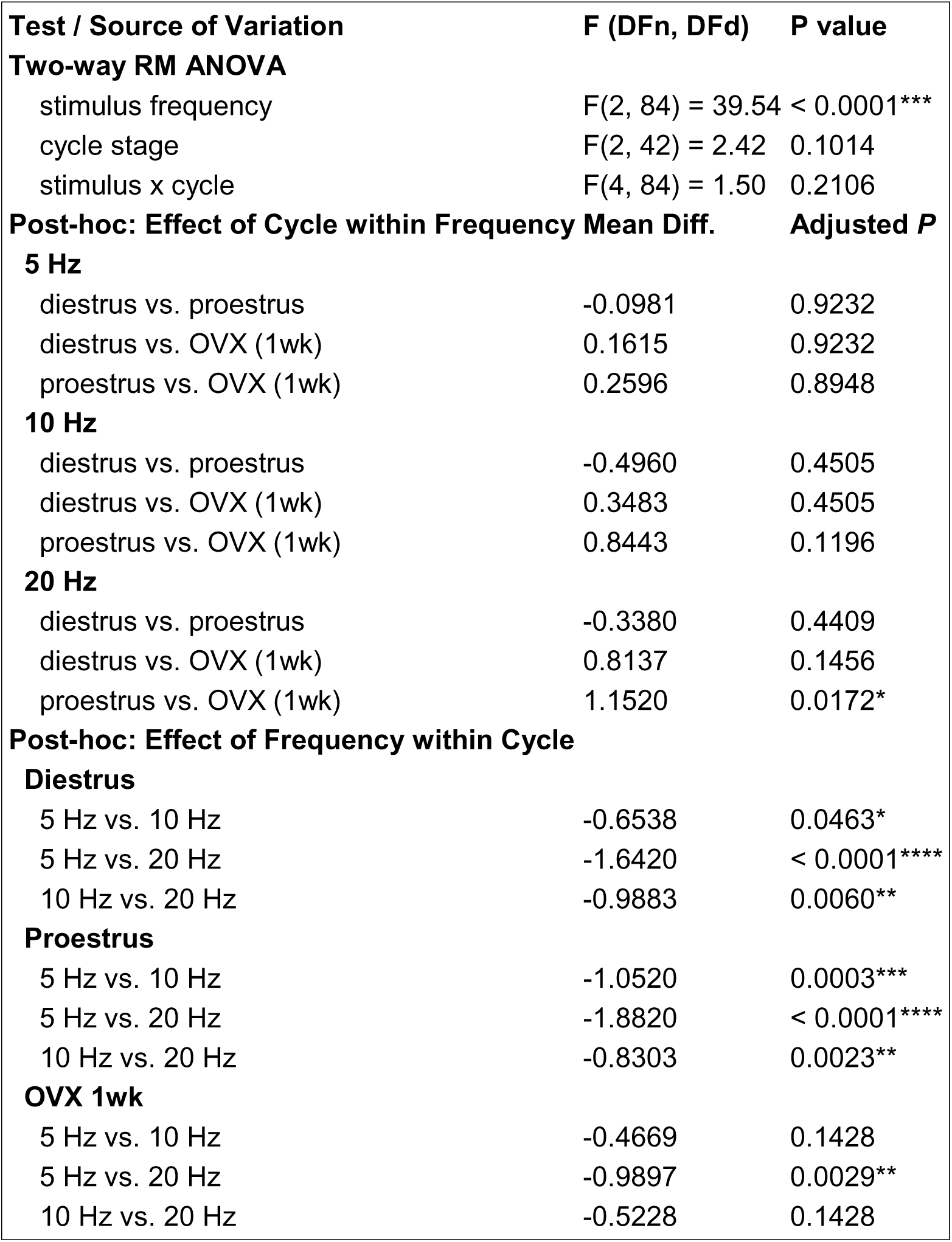
Effects of stimulation frequency and hormone state (diestrous, proestrous, OVX [1wk]) on change in firing rate.

At the end of the whole-cell recordings, some cells were given a 20Hz, 30s stimulus train to assess PSCs during high frequency stimulation (Figure 5F). Both PSC frequency (Figure 5G) and total charge transfer (Figure 5H) were greater during the stimulus than the 30s before or after the stimulus. Interestingly, there was a carryover effect of stimulus when treatment groups were combined so that charge transfer during the poststimulus period was greater than during the prestimulus period, perhaps indicating local circuit interactions persisted. These findings were consistent whether all cells or only cells with a detected synapse were considered (2-way, repeated-measures ANOVA, Table 9), indicating 20Hz stimulation of AVPV kisspeptin neurons increases PSC frequency and charge transfer at GnRH neurons, although it should be pointed out these differences are driven by substantial changes in a subpopulation of cells.

**Table 9.** Two-way ANOVA of PSC frequency and charge for 20 Hz stimulation during whole-cell recordings.

| Analysis / Source of Variation | F (DFn, DFd) or t | df | Adjusted p value |
| --- | --- | --- | --- |
| **I. PSC Frequency: All Cells** |  |  |  |
| stimulus period | F = 9.68 | 2, 92 | 0.0002*** |
| cycle | F = 1.09 | 2, 46 | 0.3458 |
| stimulus period x cycle | F = 1.21 | 4, 92 | 0.3128 |
| *Post-hoc*: pre vs. stimulus | t = 4.05 | 92 | 0.0003*** |
| *Post-hoc*: stimulus vs. post | t = 3.51 | 92 | 0.0014** |
| *Post-hoc*: pre vs. post | t = 0.54 | 92 | 0.5917 |
| **II. PSC Frequency: Synapse Only** |  |  |  |
| stimulus period | F = 10.00 | 2, 38 | 0.0003*** |
| cycle | F = 0.76 | 2, 19 | 0.4832 |
| stimulus period x cycle | F = 0.64 | 4, 38 | 0.6404 |
| *Post-hoc*: pre vs. stimulus | t = 4.16 | 38 | 0.0005*** |
| *Post-hoc*: stimulus vs. post | t = 3.51 | 38 | 0.0024** |
| *Post-hoc*: pre vs. post | t = 0.65 | 38 | 0.5168 |
| **III. PSC Charge: All Cells** |  |  |  |
| time | F = 8.49 | 1.03, 47.3 | 0.0051** |
| cycle | F = 1.21 | 2, 46 | 0.3077 |
| time x cycle | F = 1.44 | 2.06, 47.3 | 0.2479 |
| *Post-hoc*: pre vs. stimulus | t = 2.86 | 48 | 0.0126* |
| *Post-hoc*: stimulus vs. post | t = 2.39 | 48 | 0.0208* |
| *Post-hoc*: pre vs. post | t = 4.27 | 48 | 0.0003*** |
| **IV. PSC Charge: Synapse Only** |  |  |  |
| stimulus period | F = 7.96 | 1.02, 19.3 | 0.0105* |
| cycle | F = 0.55 | 2, 19 | 0.5848 |
| stimulus period x cycle | F = 0.62 | 2.04, 19.3 | 0.5528 |
| *Post-hoc*: pre vs. stimulus | t = 2.97 | 21 | 0.0219* |
| *Post-hoc*: stimulus vs. post | t = 2.88 | 21 | 0.0219* |
| *Post-hoc*: pre vs. post | t = 2.44 | 21 | 0.0235* |

### Detection of a synapse is associated with increased firing response magnitude in cells from proestrous mice

Sufficient cells were recorded at 20Hz stimulus to examine the effect of a detectable synapse on the response. This population of cells includes all the cells in Figure 5 plus additional cells in which only a 20Hz stimulus was successfully applied. Figure 6B shows firing rate before and after the stimulus train for each cell, color coded by firing response and synapse detection. The percentage of cells with synapses detected remained under 33% and was not different among groups (Figure 6C, X^2^ 1.425, df=2, p=0.4904) with most of the cells exhibiting an increase in firing rate not having a detectable synapse. Detection of a synapse was not associated with differences in baseline firing rate before the 20Hz stimulus among groups (Figure 6D, two-way ANOVA, Table 10). In contrast, the firing rate after the 20Hz stimulus was higher in cells from proestrous mice with a synapse detected than those without, and also higher than that in cells from either diestrous or OVX mice with synapses detected (Figure 6E, Table 10). Similarly, the change in firing rate in cells from proestrous mice in which a synapse was detected was greater than that in those without a synapse detected, and greater than cells from OVX mice with a synapse detected (Figure 6F, Table 10).

**Figure 6.**
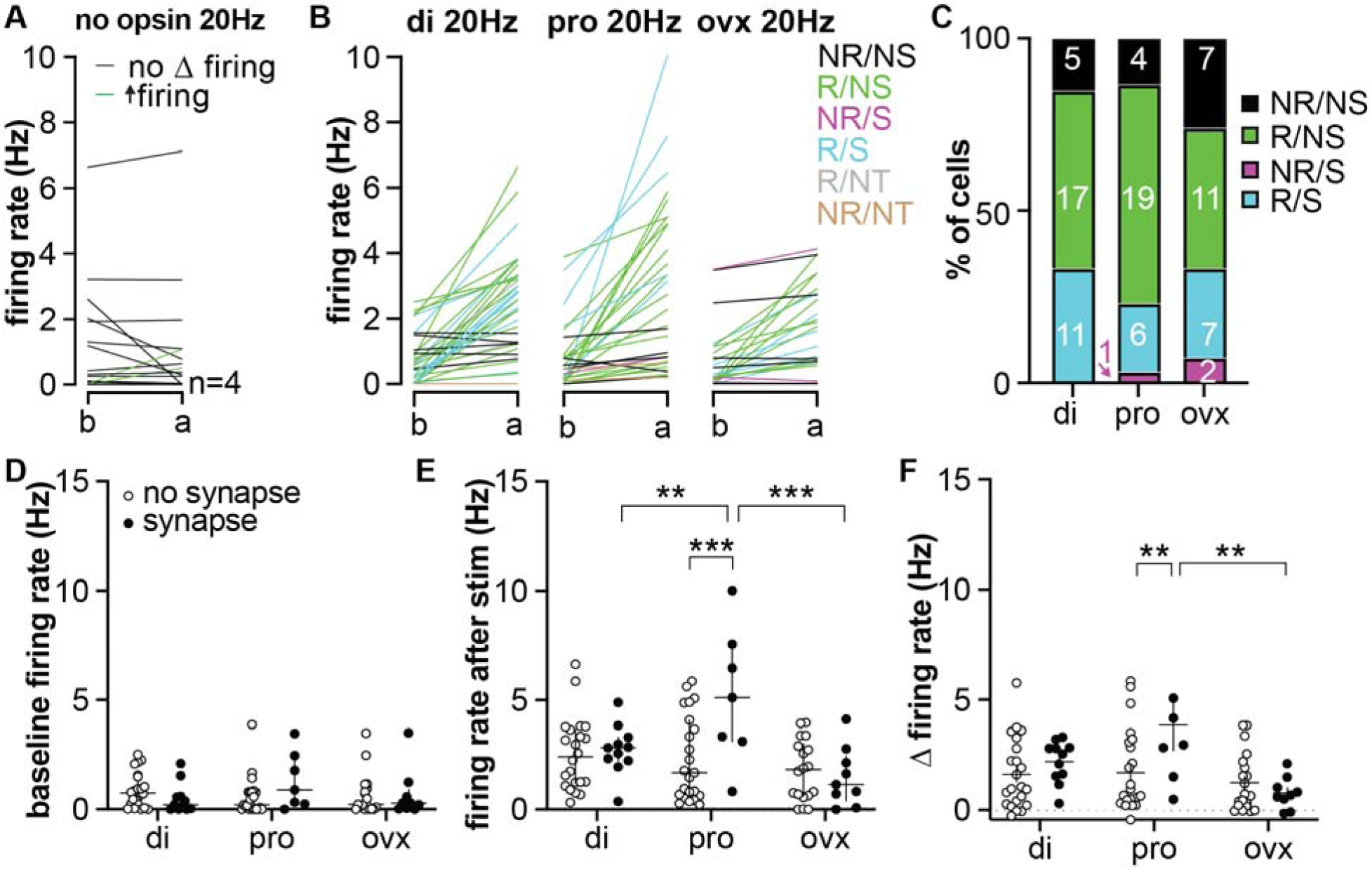
Detection of a synapse is associated with increased firing response on proestrus. A. Most GnRH neurons do not respond to 20Hz photostimulus when no opsin is present. B. Individual values for mean firing rate before (b) and after (a) 20Hz stimulus in the three groups; black non-responding cells without synapse, green responding cells without synapse, magenta non-responding cells with synapse, cyan responding cells with synapse, grey responding cells synapse not tested, non-responding cells synapse not tested. Group is indicated at top. C. Percent of cells in each category by group, cells without synapse test are omitted. Numbers in boxes are numbers of cells. D-F. Individual values and mean±SEM baseline firing rate (D), poststimulus firing rate (E), and change in firing rate (Δ, F). * *P*<0.05, ** *P*<0.01, *** *P*<0.001two-way repeated-measures ANOVA/Holm-Sidak. di, diestrus; pro, proestrus; ovx, ovariectomized.

**Table 10.** Two way ANOVA of baseline firing rate and change in firing rate in response to 20 Hz, influence of synapse.

| Test / Source of Variation | F (DFn, DFd) | P value |
| --- | --- | --- |
| **I. Baseline Firing Rate** |  |  |
| cycle stage (di, pro, OVX) | F(2, 84) = 0.60 | 0.5503 |
| synapse | F(1, 84) = 0.33 | 0.5658 |
| cycle stage x synapse | F(2, 84) = 2.28 | 0.1083 |
| **II. Stimulated Firing Rate** |  |  |
| cycle stage (di, pro, OVX) | F(2, 84) = 8.28 | 0.0005*** |
| synapse | F(1, 84) = 5.25 | 0.0245* |
| cycle x synapse | F(2, 84) = 5.38 | 0.0063** |
| **Post-hoc: Cycle Effect** | **Mean Diff.** | **Adjusted P** |
| **No Synapse** |  |  |
| diestrus vs. proestrus | 0.2302 | 0.6615 |
| diestrus vs. OVX | 0.7654 | 0.4369 |
| proestrus vs. OVX | 0.5351 | 0.5591 |
| **Synapse** |  |  |
| diestrus vs. proestrus | -2.4630 | 0.0095** |
| diestrus vs. OVX | 1.2490 | 0.1175 |
| proestrus vs. OVX | 3.7130 | 0.0002*** |
| **Post-hoc: Synapse Effect** |  |  |
| **diestrus:** no synapse vs. synapse | -0.2018 | 0.7566 |
| **proestrus:** no synapse vs. synapse | -2.8950 | 0.0003*** |
| **OVX:** no synapse vs. synapse | 0.2819 | 0.6953 |

### ePSCs are monosynaptic and GABAergic

Pharmacology experiments were done in a subset of cells to test if synapses were monosynaptic and to identify the mediating transmitter. A total of eight cells were tested with a drug application order of CNQX, TTX, 4AP, gabazine (Figure 7A) and success rates compared to assess drug effect. Not all cells received all treatments and cells with skipped treatments are shown in dashed lines in Figure 7B. ePSCs were eliminated by TTX and rescued by 4AP in all cells tested (n=5) indicating they were monosynaptic (Table 11, Figure 7B). Consistent with this, the latency for ePSCs in cells from both the daily surge (Figure 4F) and cycle models (Figure 7D) had medians between 5 and 6 ms. Success rate (proportion of stimulus pulses that evoked PSCs) was also used to identify the mediator of the current. PSCs were blocked by gabazine but not CNQX (Table 11). Partial recovery after a 10min GABAzine washout period was observed in four cells. Drug application for these studies precluded using brain slices for additional recordings. We thus used PSC latency and jitter measurements from pharmacologically-confirmed monosynaptic PSCs to identify ePSCs in drug-free recordings made in the daily surge (Figure 4) or cycle models (Figure 7) as evoked or not-evoked. In the cycle model, there were no differences among groups for ePSC success rate, latency, jitter or amplitude ratio (Table 12).

**Figure 7.**
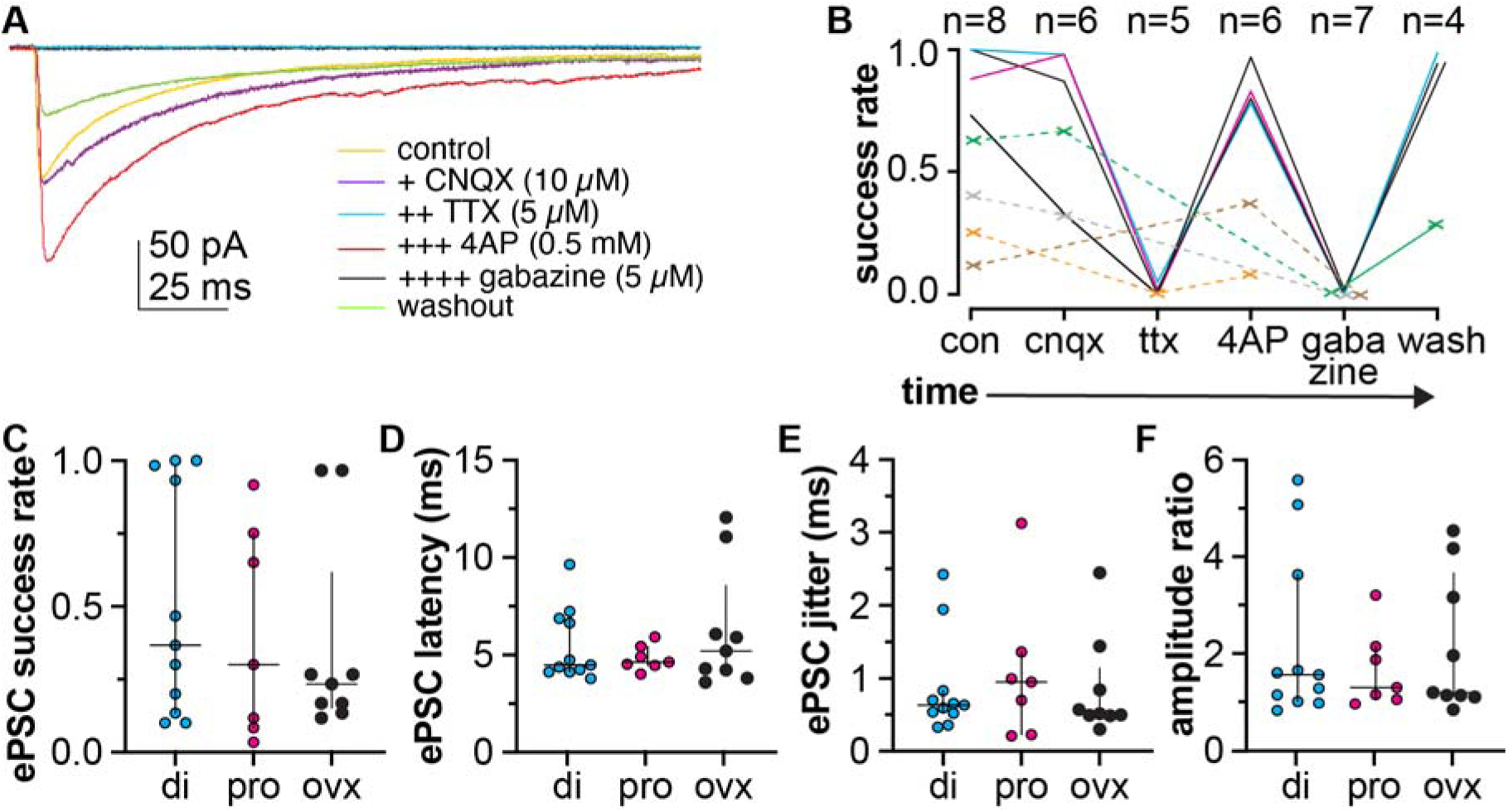
Pharmacological characterization of evoked postsynaptic currents (ePSCs) and characteristics in cells from Experiment 3. A. Mean evoked PSC traces under different pharmacological treatments given in order from top to bottom of legend. Colors in A refer to different treatments in the cell shown. B. Colors in B refer to the success rate for evoking PSCs under different conditions from the same cell. Cells with each treatment are shown as solid lines (magenta cell was lost before wash). Cells for which some treatments are skipped are shown by dashed lines with X marking the treatments given. Number of cells receiving each treatment is shown at the top of B. C-F. Individual values and median±IQR for ePSCs in proestrus (pro), diestrus (di), and ovariectomized mice 1wk post surgery (ovx).

**Table 11.**
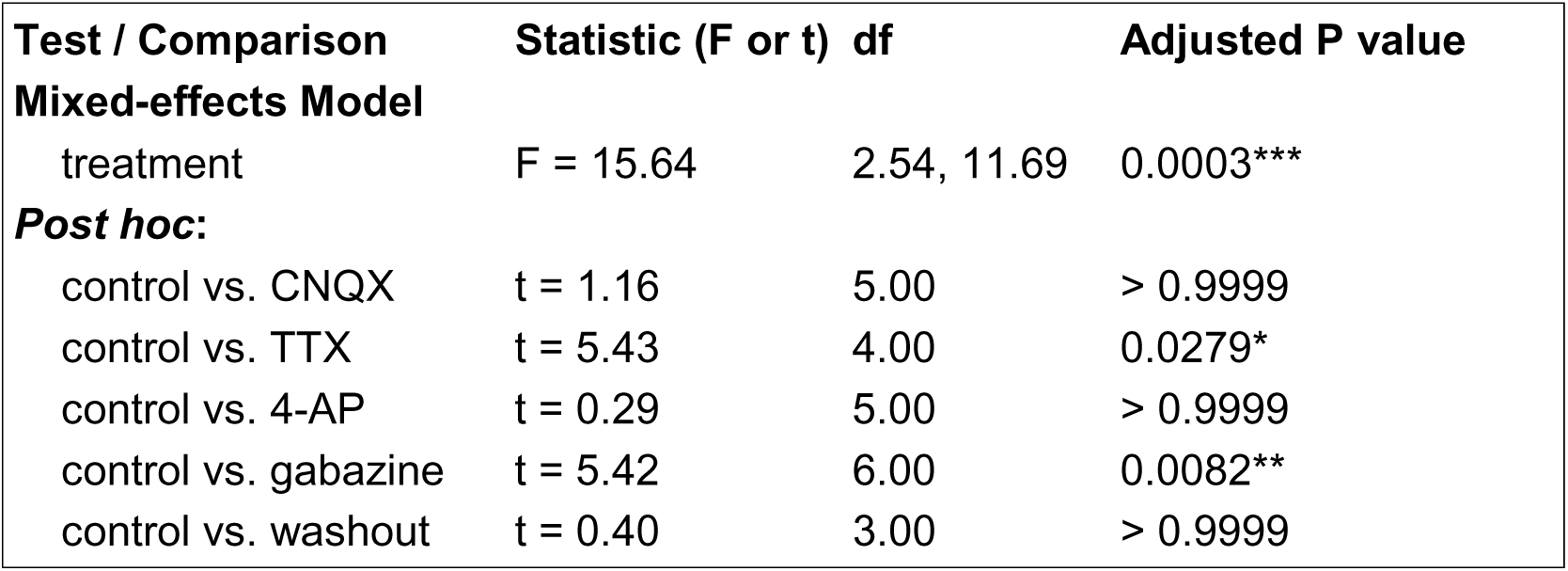
Mixed-effects analysis of pharmacological effects on PSC success rate.

**Table 12.** Kruskal-Wallis analysis of evoked PSC characteristics in cells from diestrous, proestrous, and OVX (1wk) mice.

| **Measurement** | **Diestrus**<br>**Median (IQR)** | **Proestrus**<br>**Median (IQR)** | **1wk OVX**<br>**Median (IQR)** | **Statistic (H)** | **df** | **P value** |
| --- | --- | --- | --- | --- | --- | --- |
| **success rate (%)** | 0.367 (0.85) | 0.300 (0.667) | 0.233 (0.467) | 1.26 | 2 | 0.5329 |
| **latency (ms)** | 4.516 (2.764) | 4.671 (0.988) | 5.247 (4.571) | 0.04 | 2 | 0.9787 |
| **jitter (ms)** | 0.6344 (0.310) | 0.953 (1.138) | 0.518 (0.653) | 0.58 | 2 | 0.7489 |
| **amplitude ratio** | 1.55 (2.614) | 1.286 (1.096) | 1.178 (2.546) | 0.05 | 2 | 0.9748 |

### Long-term OVX does not alter the response of GnRH neurons to optogenetic stimulation

Kisspeptin mRNA levels in the AVPV are reduced within one week of ovariectomy, but the number of neurons immunoreactive for kisspeptin peptide is not lower until six weeks post OVX (Smith et al., 2005; Brock and Bakker, 2013). To test if this decrease in kisspeptin immunoreactive cells would alter the response to photostimulation, we performed extracellular recordings on mice 6wks after OVX. Because the OVX-induced drop in kisspeptin gene transcription would affect *Cre* recombinase expression driven by that promoter, mice underwent viral injection one week before OVX, followed five weeks later by OVX ± E replacement and then extracellular recording one week later (n=6 each, Figure 8A). Groups of OVX (n=6) and OVX+E (n=4) mice were similarly prepared without viral injection for RT-PCR assessment of AVPV tissue to test the efficacy of estradiol treatment on *Kiss1* mRNA expression. Estradiol replacement elevated uterine mass in mice used for gene expression (Figure 8B, t=3.902, df-5.332, p=0.0100, Welch’s unpaired *t* test) and for recording (Figure 8D, t=5.062, df=3.005, p=0.0148, Welch’s unpaired *t* test) studies. *Kiss1* mRNA was markedly lower (Figure 8C, t=6.104, df=3.021, p=0.0087, Welch’s unpaired *t* test) in mice receiving vehicle vs estradiol implants one week before tissue collection consistent with prior work (Smith *et al., 2005)*. Cells were assessed to be photostimulus-responsive or not before identifying cells as from OVX (n=9) or OVX+E (n=10) mice. Permutation analysis revealed 3 of 9 cells from OVX mice and 3 of 10 cells from OVX+E mice responded with increased firing; an additional cell from an OVX mouse increased firing rate >2 min post onset of the stimulus and was included as a responder. Chi square found no difference (p=0.5146) between the proportion of responsive cells in the OVX and OVX+E groups. Representative extracellular recordings of a GnRH neuron not responding to photostimulation (top) and one responding (bottom) are shown in Figure 8E; both cells are from OVX+E animals. There was no difference in baseline firing rate (p=0.2110, U=29, two-tailed Mann-Whitney U test) or post-stimulus firing rate (t=1.348, df=13.60, p=0.1996, two-tailed Welch’s unpaired t test) between OVX and OVX+E groups (Figure 8F). Consistent with these observations, there was no difference in the change in firing rate from the 4 min before to the 4 min after photostimulus (t=0.3493, df=12.12, p=0.7329, two-tailed Welch’s unpaired t test, Figure 8).

**Figure 8.**
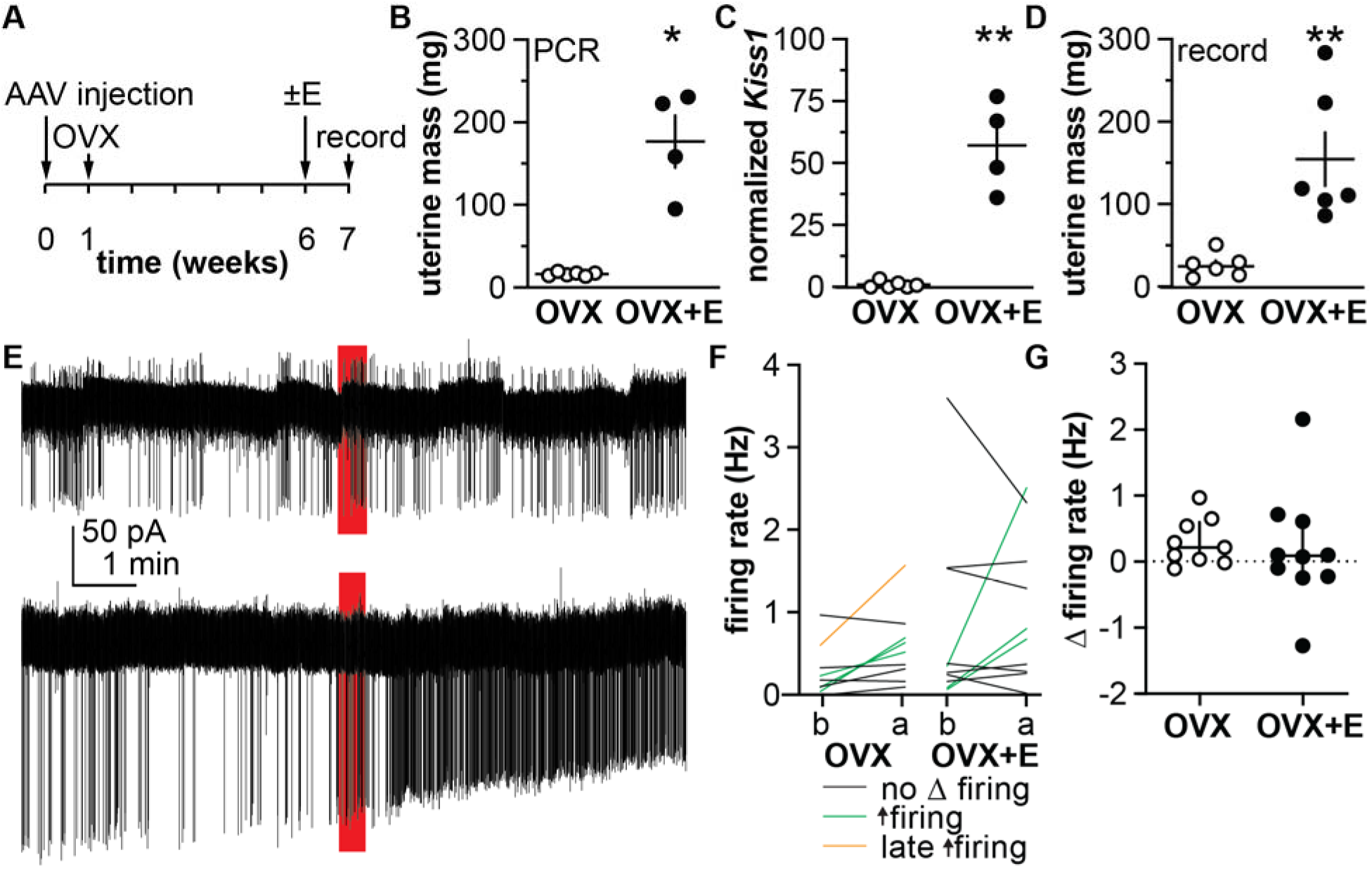
Long-term OVX does not affect GnRH neuron response to AVPV kisspeptin neuron activation. A. Experimental design. B. Individual values and mean±SEM uterine mass in mice used for PCR analysis. C. *Kiss1* expression normalized to *ActB/Rsp29*. D. Individual values and mean±SEM uterine mass in mice used for recordings. E. Representative examples of five min of recording before and after photostimulation (red bar) in a non-responding cell (top) and responding cell (bottom). F. Individual values for mean firing rate before (b) and after (a) each stimulus; green indicates responding cells, black are non-responding cells, orange is a late responding cell. G. Individual values and mean±SEM change in firing rate from before to after stimulus.

### Most GnRH neurons do not exhibit a prompt increase in firing rate upon optogenetic stimulus

Most GnRH neurons did not exhibit an immediate increase in firing rate at or near the onset of optogenetic stimulus. In addition to the before and after firing rate assessment made using the permutation analyses above, we examined firing rate at the onset of the stimulus windows. We compared the 5s just before the stimulus with the first 5s of the stimulus. Thirteen cells had an increase in firing rate of >1Hz within 5s of stimulus onset that was atypical for the firing pattern of that cell (4 diestrus, 3 proestrus, 1 OVX 1wk, 1 OVX 6wk, 1 OVX AM, 2 OVX+E AM, 1 OVX+E PM). Presence/absence of a synapse was tested in 10 of these cells and 100% of them exhibited evoked synaptic currents consistent with a monosynaptic input. A persistent increase in firing rate occurred in 73% of these cells (10 of 11 from Experiment 3, 1 of 4 from Experiment 2).

## Discussion

The preovulatory GnRH surge involves estradiol-sensitive AVPV neurons (Herbison, 2008; Uenoyama et al., 2021; Kauffman, 2022; Starrett and Moenter, 2023). GnRH neuron recordings were coupled with AVPV kisspeptin neuron fiber photostimulation to assess contributions of neuromodulators and fast-synaptic transmission. GnRH neuron firing response was modulated by hormone milieu, but monosynaptic connectivity was, unexpectedly, low and stable across hormone states tested. Evidence of monosynaptic connectivity was not required for photostimulation-induced changes in GnRH neuron firing rate, indicating a role for non-synaptic modalities.

An unbiased permutation analyses combined with a strict definition of rapid response showed increased GnRH neuron firing rate upon AVPV kisspeptin neuron photoactivation occurred on two timescales. In most cells, photostimulation induced a sustained increase in firing rate within 30-90s. A few GnRH neurons exhibited large near-instantaneous increase in firing rate upon photostimulus onset that either ceased at stimulus cessation or merged with a sustained increase in firing rate. Similar rapid and delayed-onset firing responses were reported using non-specific electrical stimulation of the AVPV and kisspeptin-targeted optogenetics (Liu et al., 2011; Qiu et al., 2016; Piet et al., 2018). More precise comparison of the time courses of these two responses to the present work is not possible due to differences in analysis approaches; the responses, however, look similar. Sustained responses were reportedly reduced by kisspeptin antagonists and were mostly absent in kisspeptin-receptor-null mice (Liu et al., 2011; Qiu et al., 2016), whereas rapid responses were reduced by blocking GABA_A_ receptors (Piet et al., 2018). In further agreement with the delayed response being neuromodulatory, we confirmed increasing the frequency of stimulation increased the prevalence and magnitude of GnRH neuron firing rate responses. This positive correlation is consistent with measurements of stimulus-secretion relationships in other neuroendocrine systems (Dutton and Dyball, 1979; Bicknell, 1988), and GnRH neurons (Chen and Moenter, 2023).

The present study extends prior work by comparing responses across different endocrine milieux and evaluating neuromodulatory and fast-synaptic systems in the same GnRH neuron. This revealed multiple important findings. First, there was a similar relationship between stimulation frequency and firing rate increase in cells from diestrous and proestrous mice, but responses were blunted 1wk post OVX. This suggests ovarian steroids help maintain the neuromodulatory signaling mode. Second, monosynaptic connectivity from AVPV kisspeptin to GnRH neurons was low and similar in the conditions tested. While low perisomatic connectivity rates are predicted by electron microscopy studies of GnRH neurons, relatively stable connectivity across hormone conditions was surprising because ovarian steroids regulate connectivity in other brain regions (Kozlowski et al., 1980; Witkin and Silverman, 1985; Witkin, 1987; Romero et al., 1994; Woolley, 1998; Lehman and Silverman, 1988). Confocal assessment of connectivity between rostral kisspeptin and GnRH neurons in mice did not observe a change in percent of GnRH neurons with appositions or number of appositions per GnRH neuron in OVX vs OVX+E mice (Kalló et al., 2012). In the present work, non-significant trends suggested lower connectivity two days vs one week after OVX. Because detected synapses between these cell types were rare, identifying a hormone effect would require an unfeasibly large sample size for adequate power. Third, a detected synapse was not required for a neuromodulatory response to occur, indicating kisspeptin signaling can occur via volume transmission and is not restricted to GnRH neurons with rare monosynaptic contacts. Finally, although not required, a monosynaptic connection was associated with larger increases in firing rate in cells from proestrous mice, suggesting the preovulatory endocrine milieu can enhance AVPV activation of GnRH neurons.

Pharmacological blockade of evoked PSCs demonstrated GABA mediates monosynaptic transmission between AVPV kisspeptin and GnRH neurons. AVPV kisspeptin neurons are GABAergic, GABA can excite GnRH neurons, and blocking GABA_A_ receptors blocks the early firing response (Cravo et al., 2011; Herbison and Moenter, 2011; Piet et al., 2018). GABAergic transmission to GnRH neurons increases during natural proestrous and hormone-induced surges (Christian and Moenter, 2007; Adams et al., 2018). AVPV kisspeptin neurons exhibit higher firing rates and increased Fos expression during surges (Adachi et al., 2007; Gusmao et al., 2022), and are a likely source of at least some increased GABAergic drive to GnRH neurons, along with lateral septal kisspeptin and the suprachiasmatic nucleus neurons (Van Der Beek et al., 1997; Szentkirályi-Tóth et al., 2025). Interestingly, a monosynaptic connection with AVPV kisspeptin neurons was detected in 8-33% of GnRH neurons but an immediate increase in firing was observed in only 6.7% (13/193). This may be attributed to the low success rate of stimulus pulses in evoking PSCs (∼2/3 of cells were near or below 50%). Low success rates were unlikely to be due to low stimulus efficacy since spike fidelity in control experiments was 100%. Rather, this suggests a lower conversion from AVPV kisspeptin neuron action potentials to a GABA release that is effective at the GnRH neuron. It is also possible that intrinsic excitability of some GnRH neurons was too low for evoked synaptic currents to induce action potentials.

In contrast to immediate responses, delayed neuromodulatory responses were common and produced sustained increases in firing rate consistent with activation of second messenger systems. We wanted to determine the mediator of this response, but were not able to block firing responses of GnRH neurons to applied kisspeptin in pilot studies, a difficulty encountered by others (Albers-Wolthers et al., 2017; Piet et al., 2018). As an alternative but indirect approach, we used mice six weeks after OVX, which reduces kisspeptin immunoreactive counts to 20% of that on proestrus (Brock and Bakker, 2013). Only about one-third of GnRH neurons from mice 6wk post ovariectomy increased firing rate upon 20Hz stimulation of AVPV kisspeptin neurons versus over two-thirds of cells from proestrous mice, possibly because of the reported loss in kisspeptin immunoreactivity. Interestingly, cells from 6wk OVX+E mice, which had restored Kiss1 mRNA expression, responded similarly to cells from 6wk OVX mice. This may indicate slower kisspeptin protein recovery following estradiol treatment, or that long-term absence of ovarian hormones causes persistent changes in the system that are less reversible. Longer-term changes to GnRH neuron excitability could also account for reduced percent of neurons responding with increased firing rate (Adams et al., 2018, 2019).

The current work has several caveats. There was an unexpected lack of detection of a positive feedback response in the daily surge model. These mice had an unmistakable increase in uterine mass, demonstrating efficacy of estradiol treatment, but the only neurobiological difference observed between groups was an increased amplitude ratio of ePSCs. While this is consistent with an increase in sPSC amplitude during the LH surge (Christian and Moenter, 2007; Adams et al., 2018), the neurobiological effects of the estradiol treatment were likely not complete at the time of study. Many GnRH neurons from daily surge mice did not exhibit a change in firing rate following AVPV kisspeptin neuron photostimulation. It is unlikely low responses were due to kisspeptin depletion as levels remain stable even at three weeks post ovariectomy (Brock and Bakker, 2013). Low rate of response may be in part attributed to the shorter time between photostimulations. There are also caveats from use of physiologic chloride levels in the pipette solution. In the first study, we attributed the low detection of synapses to reduced chloride driving force, but subsequent experiments were consistent with this low rate. Experiment 1 has low and different group sizes; it is included to avoid the impression that some groups in the daily surge model were devoid of detected synapse connections. Of note, in whole-cell recordings it is possible synapses distal to the pipette are not detected because of technical limitations (Spruston et al., 1993). Similar somatic recordings of spontaneous GABAergic PSCs in GnRH neurons, however, rarely fail to detect synaptic inputs, suggesting that some GABAergic inputs to GnRH neurons could originate from non-AVPV kisspeptin sources (Sullivan et al., 2003; Christian and Moenter, 2007; Farkas et al., 2010; Liu et al., 2017; Adams et al., 2018).

AVPV kisspeptin neurons activate GnRH neurons through two signaling modes: fast-synaptic GABAergic transmission and slow neuromodulatory signaling likely via kisspeptin. The discovery that direct synaptic connections are rare and not required for neuromodulatory responses indicates kisspeptin can act via volume transmission. AVPV kisspeptin neurons could thus activate a spatially-diffuse GnRH network despite limited monosynaptic connectivity. Ovarian steroids play a mainly permissive role by maintaining the efficacy of neuromodulatory signaling, except on proestrus, when GnRH neurons receiving monosynaptic signaling from AVPV kisspeptin neurons exhibit enhanced neuromodulatory responses, possibly serving as a hub for surge initiation. Preovulatory surge initiation is likely the result of concurrent shifts at multiple levels including increased excitability and activation of the AVPV kisspeptin neuron population (Smith et al., 2005; Adachi et al., 2007; Piet et al., 2013; Zhang et al., 2015; Wang et al., 2016; Starrett et al., 2021; Gusmao et al., 2022), strengthened functional connectivity with a subset of GnRH neurons, increased GnRH neuron excitability (Liu and Herbison, 2011; Adams et al., 2018, 2019), enhanced GnRH release (Moenter et al., 1991; Glanowska et al., 2012), and increased pituitary sensitivity to GnRH (Clarke and Cummins, 1984; Silveira et al., 2017).

## Acknowledgements

We thank Laura Burger, Elizabeth Wagenmaker, R. Anthony DeFazio, and Xi Chen for expert technical assistance. We also thank James L. Kenyon, University of Nevada, Reno, for the Excel spreadsheet used to calculate junction potentials.

## Abbreviations

GnRH: gonadotropin-releasing hormone
LH: luteinizing hormone
ERα: estrogen receptor alpha
AVPV: anteroventral periventricular area
POA: preoptic area
GFP: green fluorescent protein
Cre: Cre recombinase
AAV: adeno-associated virus
OVX: ovariectomized
OVX+E: ovariectomized plus estradiol implant
E: estradiol
kiss: kisspeptin
LED: light-emitting diode
FWHM: full width at half maximum
ACSF: artificial cerebrospinal fluid
TTX: tetrodotoxin
4-AP: 4-aminopyridine
CNQX: 6-cyano-7-nitroquinoxaline-2,3-dione
PSC: postsynaptic current
ZT: Zeitgeber time
ePSC: evoked post-synaptic current

